# Selection leads to remarkable variability in the outcomes of hybridization across replicate hybrid zones

**DOI:** 10.1101/2022.09.23.509250

**Authors:** S. Eryn McFarlane, Joshua P. Jahner, Dorothea Lindtke, C. Alex Buerkle, Elizabeth G. Mandeville

## Abstract

Hybrid zones have been viewed as an opportunity to see speciation in action. When hybrid zones are replicated, it is assumed that if the same genetic incompatibilities are maintaining reproductive isolation across all instances of secondary contact, those incompatibilities should be identifiable by consistent patterns in the genome. In contrast, changes in allele frequencies due to genetic drift should be idiosyncratic for each hybrid zone. To test this assumption, we simulated 20 replicates of each of 12 hybrid zone scenarios with varied genetic incompatibilities, rates of migration, selection and different starting population size ratios of parental species. We found remarkable variability in the outcomes of hybridization in replicate hybrid zones, particularly with Bateson-Dobzhansky-Muller incompatibilities and strong selection. We found substantial differences among replicates in the overall genomic composition of individuals, including admixture proportions, inter-specific ancestry complement, and number of ancestry junctions. Additionally, we found substantial variation in genomic clines among replicates at focal loci, regardless of locus-specific selection. We conclude that processes other than selection are responsible for some consistent outcomes of hybridization, whereas selection on incompatibilities can lead to genomically widespread and highly variable outcomes. We highlight the challenge of mapping between pattern and process in hybrid zones and call attention to how selection against incompatibilities will commonly lead to variable outcomes. We hope that this study informs future research on replicate hybrid zones and encourages further development of statistical techniques, theoretical models, and exploration of additional axes of variation to understand reproductive isolation.

## Introduction

*“Same thing you do when you cross an elephant and a rhino. El-eph-I-no.”* — Ted Lasso

Following and during speciation, the local or global persistence of novel species will commonly be tested through range shifts and secondary contact between diverged lineages. Species will be more likely to persist in the face of potential interspecific gene flow if hybrid fitness is low due to incompatible alleles or genotypes (Coyne & Orr 2004). For some groups, we have good descriptions of the genetics of incompatibilities and evolutionary models for their origin (Fishman & Sweigart 2018, Larson *et al*. 2018, Maheshwari & Barbash 2011, Orr 1995, Powell *et al*. 2020). Likewise, we have well-developed expressions for equilibrium clinal variation following secondary contact (Barton 1979, 1983, Barton & Hewitt 1985, Barton & Gale 1993, Endler 1977). Together these have formed an expectation that we can use empirical observations of gene exchange and ancestry variation in hybrid zones to identify exceptional loci. Specifically, we can use hybrid zones to look for genetic loci associated with reproductive isolation and variation in fitness between hybrids and parental type individuals. For example, geographic and genomic cline methods identify loci with deviations in allelic ancestry or state relative to a genome wide distribution of ancestry (Gompert & Buerkle 2012, Szymura & Barton 1986). There is growing interest in using contrasts of multiple hybrid zones to build our understanding of their consistent and idiosyncratic features (e.g. Simon *et al*. 2021, Westram *et al*. 2021). This mirrors diverse research in evolutionary biology that uses replicates or multiple instances of evolution to reveal parallel or concordant evolution and to implicate predictable and potentially deterministic features of evolution (Elmer *et al*. 2014, Ferris *et al*. 2021, Tenaillon *et al*. 2016).

In hybrid zones, outcomes of secondary contact and the composition of hybrid populations will be driven by drift, gene flow, selection, and attributes of the genome such as linkage and recombination rates. It is an open question what level of consistency among replicate hybrid populations we should expect with different levels of selection against recombinant genotypes. Would high consistency among replicates indicate strong selection and effective reproductive isolation, or would consistency among replicates instead be a hallmark of neutral, clinal variation? We might expect consistency to be indicative of natural selection, particularly directional selection (Pardo-Diaz *et al*. 2012, Whitney *et al*. 2006). Indeed, a number of studies have suggested some measure of locus-specific consistency as a way to determine candidate regions for incompatibility loci (e.g. Kingston *et al*. 2017, Gagnaire *et al*. 2013, Caballero-Villalobos *et al*. 2021, Larson *et al*. 2014). However, the fitnesses of hybrid genotypes that contribute to intrinsic post-zygotic reproductive isolation do not involve directional selection, but are predicted to be multilocus, epistatic incompatibilities (Bateson 1909, Dobzhansky 1937, Muller 1942). Rather than deterministic and consistent shifts in allele frequency, the evolutionary dynamics associated with these fitness matrices include unstable equilibria and frequency dependence that could lead to high variance in outcomes, despite strong selection.

In a world in which the evolutionary processes governing hybridization generate predictable outcomes, one could map between empirical observations and evolutionary processes. Specifically, hybrid zone characteristics (e.g. genetic composition and ancestry of individuals, number of hybrids, time since contact, and relative abundances of parental species) would allow inference of the different demographic conditions, including selection and gene flow. Moreover, replicate hybrid zones could be used to understand how these evolutionary processes vary across space, time, and ecological variation (Rieseberg *et al*. 1999, Buerkle & Rieseberg 2001, Harrison & Larson 2016, Westram *et al*. 2021). Consistency or parallelism of loci with exceptional differentiation or introgression between populations (via F_ST_, and geographic or genomic clines) has been suggested as evidence for either loci associated with adaptive introgression or genomic regions associated with reproductive isolation (Langdon *et al*. 2022, Matute *et al*. 2020, Rieseberg *et al*. 1999, Simon *et al*. 2020, 2021, Westram *et al*. 2021). Explicitly stated, this predicts that consistent selection against a hybrid genotype, regardless of environment, will lead to detectable consistency across replicate hybrid zones, and this consistency is a hallmark of selection. Further, this assumes that replicate hybrid zones between the same species pairs will behave the same way (i.e. similar to replicated lab populations) across time or space. But the question remains: if hybrid zones are indeed windows into evolution (Harrison 1990), do different scenes viewed through different windows imply differences in process? If instead the outcomes of hybridization are highly idiosyncratic or capricious (*sensu* Lewontin 1966), potentially due to stochasticity following initial contact in different hybrid zones, the ability to link patterns to processes will be limited no matter how many replicate hybrid zones are sampled.

To resolve expectations for consistency and contingency in evolutionary replicates, we simulated replicate outcomes of hybridization, varying selection, migration, ratio of parental species, and genetic architecture of incompatibilities. We have deliberately simulated only simple genetic architectures, specifically two or four locus incompatibilities, rather than tackling polygenic selection. Specifically, we simulated either an inter-genomic (hereafter ‘BDMI’, i.e. a Bateson-Dobzhansky-Muller incompatibility) or intra-genomic incompatibility (hereafter ‘pathway’). BDMIs are theorized as the result of different alleles arising at two (or more) loci in diverged lineages, where an incompatibility arises when the lineages hybridize, and heterozygous genotypes have a negative epistatic interaction, resulting in lower fitness in *F*_1_, *F*_2_, or both categories of individuals (Bateson 1909, Dobzhansky 1936, Muller 1942). For example, (Powell *et al*. 2020) found a two locus incompatibility that contributes to melanoma in *Xiphophorus maculatus* and *X. hellerii* hybrids. In contrast, pathway incompatibilities have been associated with *cis* regulation of gene expression, where incompatibilities can result in disregulated expression in hybrids (Landry *et al*. 2007).

We are particularly interested in strictly intrinsic incompatibilities (i.e., hybrids have lower fitness without ecological, or extrinsic selection (Coughlan & Matute 2020)). Intrinsic genetic incompatibilities are expected to be the final step of reproductive isolation, preventing gene flow between species even if there is no longer prezygotic isolation, such as in the case of range shifts or translocations (Coughlan & Matute 2020). There is an expectation that intrinsic incompatibilities could be more general across taxa than mechanisms leading to prezygotic isolation (Xiong & Mallet 2022). Once we understand how intrinsic incompatibilities in consistent environments might result in variable outcomes across sites, we can develop theoretical expectations for more complex processes. For example, there are not yet theoretical expectations for how ecological interactions with genetic incompatibilities (Thompson *et al*. 2022) or ‘hybridization load’ (Moran *et al*. 2021) might behave across replicate hybrid zones, or, eventually even across taxa. Theoretical understanding of the variability of intrinsic incompatibilities across replicate hybrid zones is needed as a baseline before adding the additional complications of ecological selection.

In addition to varying selection, migration, and incompatibility type, we investigated hybridization outcomes across a range of starting population size ratios of parental species, from 1:1 to 50:1. Uneven ratios of parental species may occur when a small number of long-distance migrants disperse into the range of a related species, or on range margins where one species is much more common. Anthropogenic hybridization also likely features unequal starting parental contributions, related to a history of conservation translocations or range expansions, we also wanted to ask to what extent selection could lead to consistency when there are unequal parental population sizes.

Many simulations we use here were originally performed in association with Lindtke & Buerkle (2015), but were not previously used to quantify consistency in outcomes of hybridization. Lindtke & Buerkle (2015) demonstrated that different genetic architectures causing reproductive isolation resulted in variation in introgression and the effectiveness of reproductive isolation. In contrast, we examined variation among replicates to 1) quantify the degree of variability across replicate hybrid zones with identical evolutionary conditions (genomic architecture and fitness consequences of incompatibilities, population size, migration); 2) determine how different evolutionary conditions affect consistency across replicate hybrid zones; and 3) contrast variability of genotypes at specific loci, particularly those underlying the incompatibilities between species. Finally, we examined how consistency among populations increased with unequal parental populations and to what extent this ratio between parental populations made it easier to pinpoint effects of selection on locus-specific introgression. We hypothesize that the effects of drift at both the population and locus-specific levels make it difficult to detect selection. We predict that even when there is consistency among replicate hybrid populations, as is expected when there parental population sizes are unequal, the consistency is not due to strong selection at either the individual or locus-specific level, but rather due to decreased opportunity for stochastic and highly variable outcomes of hybridization.

## Methods

### Simulation framework

Following Lindtke & Buerkle (2015) and Gompert *et al*. (2012), we simulated diploid whole genomes (2N = 20) using the software dfuse (https://www.uwyo.edu/buerkle/software/dfuse/, https://doi.org/10.5061/dryad.0506g). A total of 510 loci were simulated per individual, with 51 equally spaced loci per chromosome. Individuals were arranged in a chain of 11 demes or 3 demes, with infinitely sized populations of different species at either end of the chain of demes and 150 individuals in the intermediate demes. Thus, the meta populations act as sink populations, where there can be admixture from the infinite parental source populations (Fig. 1, Lindtke & Buerkle 2015)). Simulations of 11 demes allowed for population structure, with individuals in the spatially central deme unlikely to have direct parentage involving the two species (F1 or backcrossed hybrids). This is consistent with a ‘classical’ hybrid zone, rather than a ‘mosaic’ hybrid zone (Rand & Harrison 1989).

**Figure 1:**
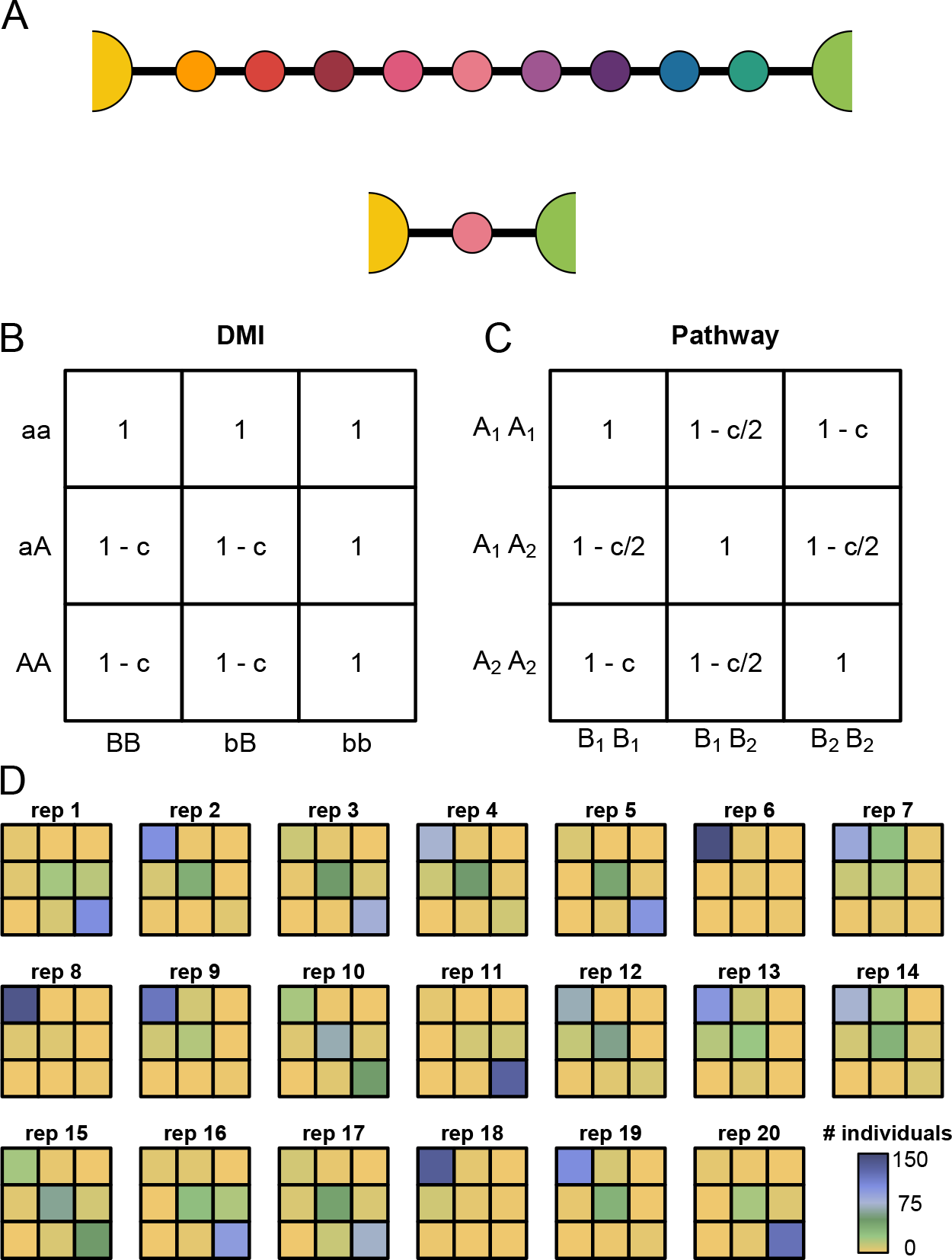
(A) Analyses were based on the outcomes of hybridization in either 11 deme or 3 deme simulations. Genotype fitness matrices are shown for the (B) BDMI and (C) pathway genetic architectures. Levels of selection (*c*) were set to either 0, 0.2, or 0.9. (D) The number of individuals from generation 10 having each genotypic combination for the two BDMI loci (1.4 and 1.6) is displayed for each of the twenty replicates from deme 6 of the BDMI, migration (*m*) = 0.01, selection (*c*) = 0.9 scenario.

To determine the degree of variability among replicate hybrid zones across a broad range of evolutionary conditions, we considered 20 independent replicates of 12 different scenarios that varied by migration rate, strength of selection, and type of incompatibility. Migration rate was set as *m/*2, where *m* was either 0.01 or 0.2 and could occur in a stepwise fashion between demes (except from deme 2 to 1 and from deme 10 to 11 (deme 2 to 3 in 3-deme simulations), where migration was not allowed). Viability selection (*c*) against individuals carrying mismatched alleles in either a BDMI or pathway incompatibility was either 0, 0.2, or 0.9. For all BDMI scenarios, two loci on the first chromosome (the fourth and sixth loci on the first chromosome: 1.4, 1.6) were specified as the causal variants, whereas two loci on both the first and second chromosomes (1.2, 1.4, 2.2, 2.4) were specified for all pathway scenarios (see Fig. 1 for genotypic fitness values). Incompatibility loci positions were chosen arbitrarily while maintaining the same distance between markers (Lindtke & Buerkle 2015).

Individual genotypes, admixture proportion (*q*), inter-specific ancestry (*Q*_12_), and the number of junctions between ancestry blocks were all recorded for each individual every 10 generations (from 10–100). The latter three statistics describe the ancestry of hybrid individuals: the admixture proportion (*q*) is the proportion of ancestry in an individual from one of the two parental species, the inter-specific ancestry (*Q*_12_) is the proportion of an individual’s diploid genome where one copy comes from parental population 1 and the other parental population 2, (Fig. S1; Gompert *et al*. 2014, Lindtke *et al*. 2014, Shastry *et al*. 2021), and the number of junctions describes the extent of recombination between ancestry from the parental genomes. With more generations of hybridization and recombination, inter-specific ancestry (*Q*_12_) declines and the number of junctions increases. We chose these metrics to describe our hybrid individuals, as the distributions of *q*, *Q*_12_, and junctions have been used to infer the processes underlying hybrid zones (Jiggins & Mallet 2000, Schumer *et al*. 2017). We also used genomic clines to describe introgression at individual loci (Gompert & Buerkle 2011), to better describe hybridization outcomes at individual loci in addition to the description of overall ancestry of hybrid individuals described by the metrics above. A more comprehensive description of the simulations can be found in Lindtke & Buerkle (2015).

### Unequal parental population sizes

Because we expected rare alleles to go to fixation quickly in small populations, we focused our simulations initially on scenarios with equal proportions of each parental type and primarily present these results. However, unequal parental contributions may be common under some scenarios, so we also simulated other starting ratios of parental population sizes. For these analyses, we used a 3 deme scenario (Fig. 1; as in Lindtke & Buerkle 2015) because this made it more intuitive to directly change the relative contributions of parental populations. Specifically, instead of specifying 150 individuals per deme, as above, we varied the population sizes in demes 1 and 3 to manipulate the parental population sizes, specifying ratios of 50:1, 10:1, 5:1, 2:1 and 1:1 (Table 1). We initially predicted that unequal parental populations could lead to less variation among replicates, as it is expected that more common alleles go to fixation according to their frequency (Kimura 1962).

**Table 1:** We varied the ratio of the starting parental population sizes at either end of three deme simulations, where we simulated BDMIs, migration rates of each 0.01, 0.2, and selection of each 0, 0.2 and 0.9. This was to test how uneven population ratios affected consistency across replicate hybrid zones in combination with varying effects of selection. We report here the range of F statistics, p values and *R*^2^ to demonstrate the similarity in results, regardless of the ratio of contribution from parental populations.

| Parental Population Sizes | Population size ($N$) | | | Admixture proportion ($q$) | | | Inter-specific ancestry ($Q_{12}$) | | | Junctions | | |
| --- | --- | --- | --- | --- | --- | --- | --- | --- | --- | --- | --- | --- |
| | Deme 1 | Deme 2 | Deme 3 | F range | p range | $R^2$ range | F range | p range | $R^2$ range | F range | p range | $R^2$ range |
| 1:1 | 150 | 150 | 150 | 1.28 – 990 | p<0.001 – 0.182 | 0.005 – 0.863 | 0.423 – 249 | p<0.001 – 0.986 | 0.002 – 0.614 | 1.24 – 214 | p<0.001 – 0.213 | 0.005 – 0.577 |
| 2:1 | 150 | 150 | 300 | 1.16 – 180 | p<0.001 – 0.283 | 0.007 – 0.534 | 0.90 – 72 | p<0.001 – 0.583 | 0.006 – 0.315 | 1.70 – 126 | p<0.001 – 0.029 | 0.11 – 0.446 |
| 5:1 | 60 | 150 | 300 | 1.97 – 481 | always p<0.001 | 0.012 – 0.754 | 0.71 – 348 | p<0.001 – 0.818 | 0.004 – 0.690 | 2.96 – 299 | always p<0.001 | 0.18 – 0.656 |
| 10:1 | 30 | 150 | 300 | 2.32 – 359 | always p<0.001 | 0.015 – 0.696 | 1.69 – 343 | p<0.001 – 0.031 | 0.011 – 0.686 | 3.92 – 289 | always p<0.001 | 0.024 – 0.648 |
| 50:1 | 6 | 150 | 300 | 2.64 – 1001 | always p<0.001 | 0.017 – 0.865 | 1.62 – 700 | p<0.001 – 0.044 | 0.010 – 0.82 | 5.31 – 384 | always p<0.001 | 0.033 – 0.710 |

### Quantifying variability across replicate hybrid zones

Our primary goal was to determine under which scenarios replicate hybrid zones are consistent with one another. For all analyses, we considered only the central deme where the proportion of hybrids is typically the highest (either deme 6 or deme 2, see Fig. 1). Additionally, we focused on the outcomes from generations 10 and 100. As our primary metric of variability, we calculated Fisher’s F statistics from analysis of variance (ANOVA) in R v4.0.5 (R Core Team 2021) to quantify the ratio of among versus within replicate variance for a given simulation scenario. Larger F statistics correspond to a greater variance among replicates relative to variation within replicates. We also report the *R*^2^, the proportion of variance explained by replicate.

We first focused on the variability of individual level parameters by calculating ANOVA F statistics based on *q*, *Q*_12_, and the number of transitions (junctions) in ancestry along chromosomes. Next, we asked to what extent specific genetic loci respond similarly across replicates. To do this, we calculated ANOVA F statistics based on the genotypes of three specific loci: a locus directly under selection (locus 1.4); a locus on the same chromosome as selected loci (locus 1.10); and a physically unlinked locus (3.4). In addition to ANOVA on genotypes, we performed ANOVA on minor ancestry. We set the response variable as either the locus specific genotype (0, 0.5 or 1), if mean *q* for the replicate was *<* 0.5, or, if *q >* 0.5 then we had the response variable as 1-genotype. This ensures that a minor ancestry score of 1 meant an individual had two copies of the rarer ancestry at the replicate level. As a complementary approach to assess genotypic variability among individuals and replicates, we used principal component analysis (PCA) in R based on raw genotypes from all 510 loci. For each replicate, we calculated a 90% confidence ellipse of the first two PCs using the *ordiellipse* function in vegan v2.6-2 (Oksanen *et al*. 2022). Finally, we examined the variability in the genetic composition of hybrids using using multinomial regression that reflects the probabilities of homozygous and heterozygous genotypes given an individual’s genome-wide admixture proportion (Gompert & Buerkle 2010). We consider any significant effect of replicate identified by ANOVA to be evidence of variability, although we recognize that statistically significant differences could arise through variation that is not large in magnitude.

## Results

We examined 12 scenarios that varied with respect to the degree of gene flow (*m* = 0.01 versus 0.2), strength of selection (*c* = 0 versus 0.2 versus 0.9), and genetic architecture (BDMI versus pathway). In general, proportionally more variation in any of the response variables (*q*, *Q*_12_, or number of junctions) was associated with differences among replicates (i.e. had larger ANOVA F statistics) when selection was high, gene flow was low, and when incompatibilities were due to a BDMI (Fig. 2). At generation 10, there was significant variation explained by replicate in number of junctions in all 12 scenarios (F range: 3.26–146, *R*^2^ range: 0.020–0.482), in admixture proportion (*q*) for all 12 scenarios (F range: 2.16–1148, *R*^2^ range: 0.014–0.880), and in inter-specific ancestry (*Q*_12_) for 12 scenarios (F range: 0.72–153, *R*^2^ range: 0.005–0.494) (Table S1). For the three deme structure, we also found substantial among replicate variation in *q* when selection was high and migration was low (Fig. S2), but the high selection, high migration scenario was more consistent than in the 11 deme structure. At generation 100, replicates differed in the number of junctions for all 12 scenarios (F range: 17.76–1183, *R*^2^ range: 0.079–0.883), in *q* for all 12 scenarios (F range: 7.24–3094, *R*^2^ range: 0.044–0.952), and in *Q*_12_ for 11 scenarios (F range: 1.07–433, *R*^2^ range: 0.007–0.734) (Table S1). For those scenarios with the largest F statistics for *q* and *Q*_12_, extreme variation was not isolated to the chromosome with incompatible loci, but instead extended throughout the genome (Fig. 3; note that *Q*_12_ was less variable in the three deme structure, Fig. S3). Accordingly, in these scenarios the genetic composition of hybrids varied greatly across replicates for key characteristics such as the proportion of F2 hybrids and the direction of backcrossing (Fig. 4), although there was less variation among replicates in the three deme case (Fig. S4). Across scenarios, log-transformed F statistics for *q* and *Q*_12_ were highly correlated with one another in generation 10 (r = 0.874; Fig. S5) and generation 100 (r = 0.933; Fig. S6). Further, a replicate’s median *q*, and to a lesser extent *Q*_12_, in generation 10 were often correlated with its median value in generation 100, especially when selection was strong (*c* = 0.9) (Figs. S7, S8).

**Figure 2:**
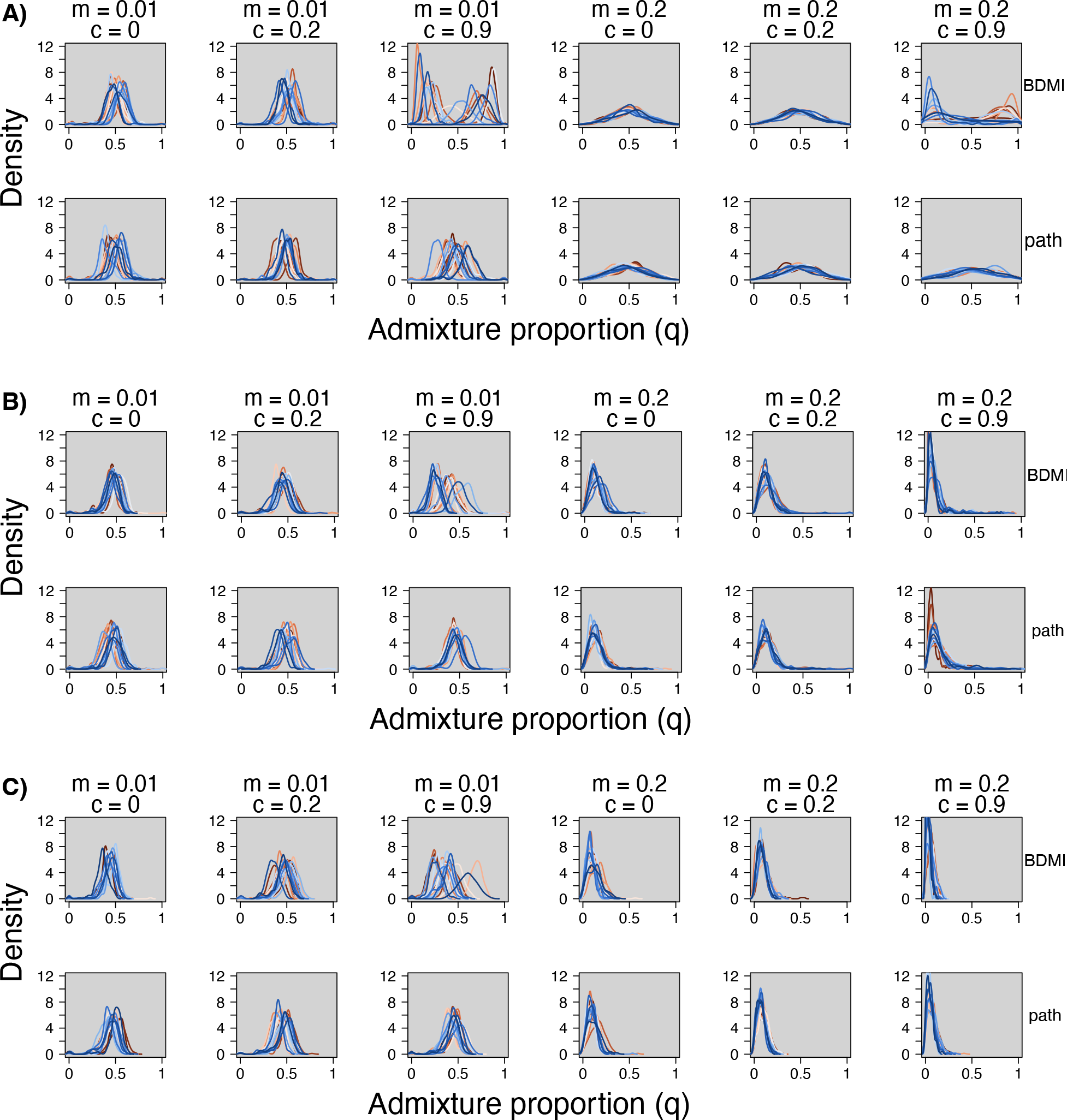
When initial parental population sizes are equal (A), selection leads to remarkable variation in the outcomes of hybridization across replicate hybrid zones. When parental species are initially at a 10:1 ratio (B) or 50:1 ratio (C), outcomes of hybridization are still variable under conditions of low migration and high selection, especially with the BDMI architecture. For all figure panels, the distribution of admixture proportion (*q*) from generation 10 is shown for 20 replicates from the 12 scenarios that varied by migration (*m*), selection (*c*) and incompatibility (BDMI versus pathway). Colors correspond to different replicates and match those in other figures. See Fig. S6 for results from generation 100 for equal parental population sizes. Panel A is from an 11 deme scenario (as are Figures 3 and 4, while Panels B and C are from 3 deme scenarios. The 3 deme scenario for even parental population sizes is Supplementary Fig. S2.

**Figure 3:**
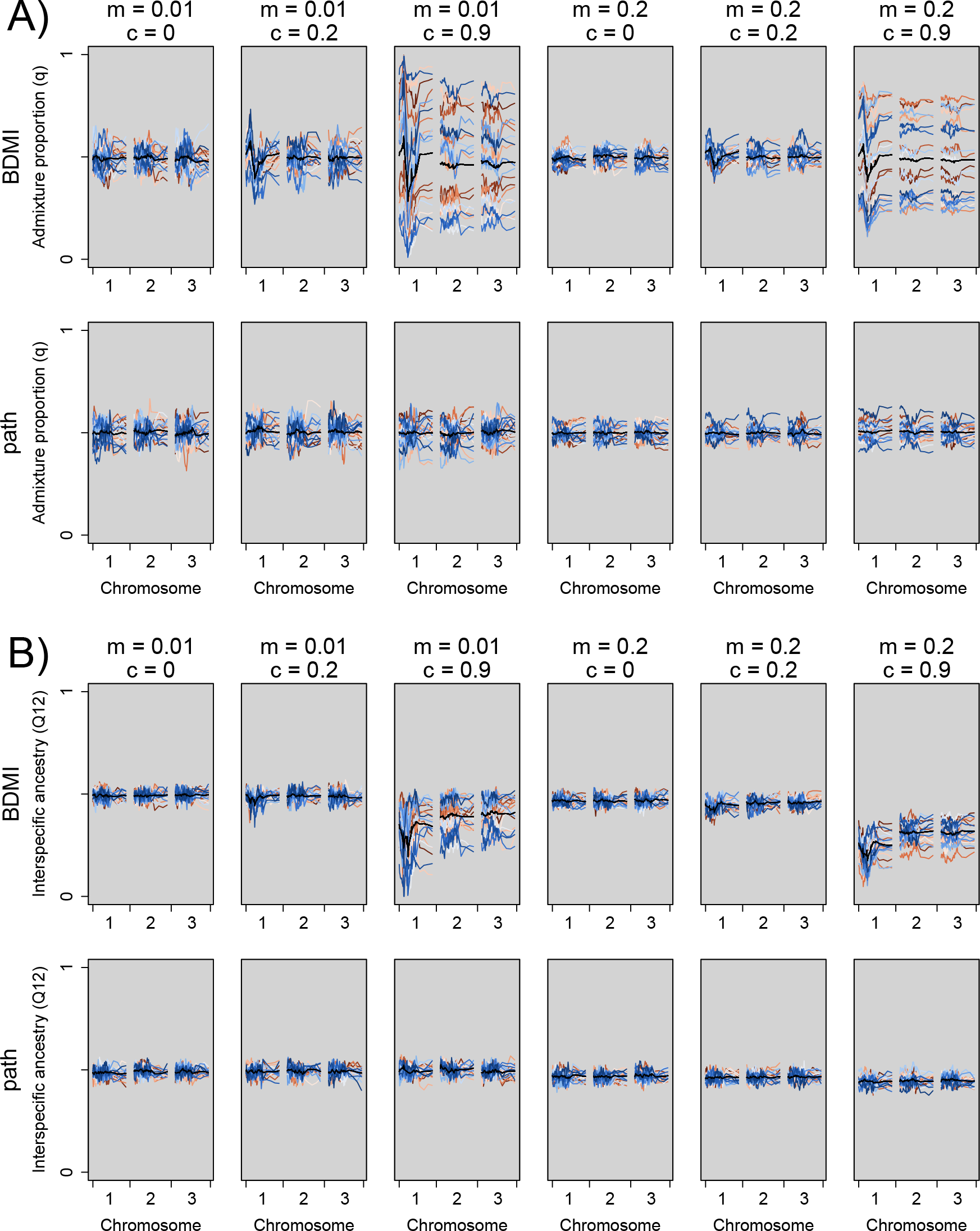
Locus-specific variation in average (among individuals in a replicate) admixture proportion (*q*, panel A) and inter-specific ancestry (*Q*_12_, panel B) for replicates (different colours), six combinations (columns) of migration (*m* = 0.01 or 0.2) and selection (*c* = 0, 0.2 or 0.9), and two types of incompatibilities (BDMI or pathways, in rows). Colors correspond to different replicates and match those in other figures. The black line is the average across all replicates, as in Lindtke & Buerkle (2015). The plots are for simulations after ten generations. Chromosome 1 has loci with BDMI or pathway incompatibilities, chromosome 2 has pathway incompatibility loci, while chromosome 3 has only neutral loci, and is representative of the rest of the genome (chromosomes 4–10).

**Figure 4:**
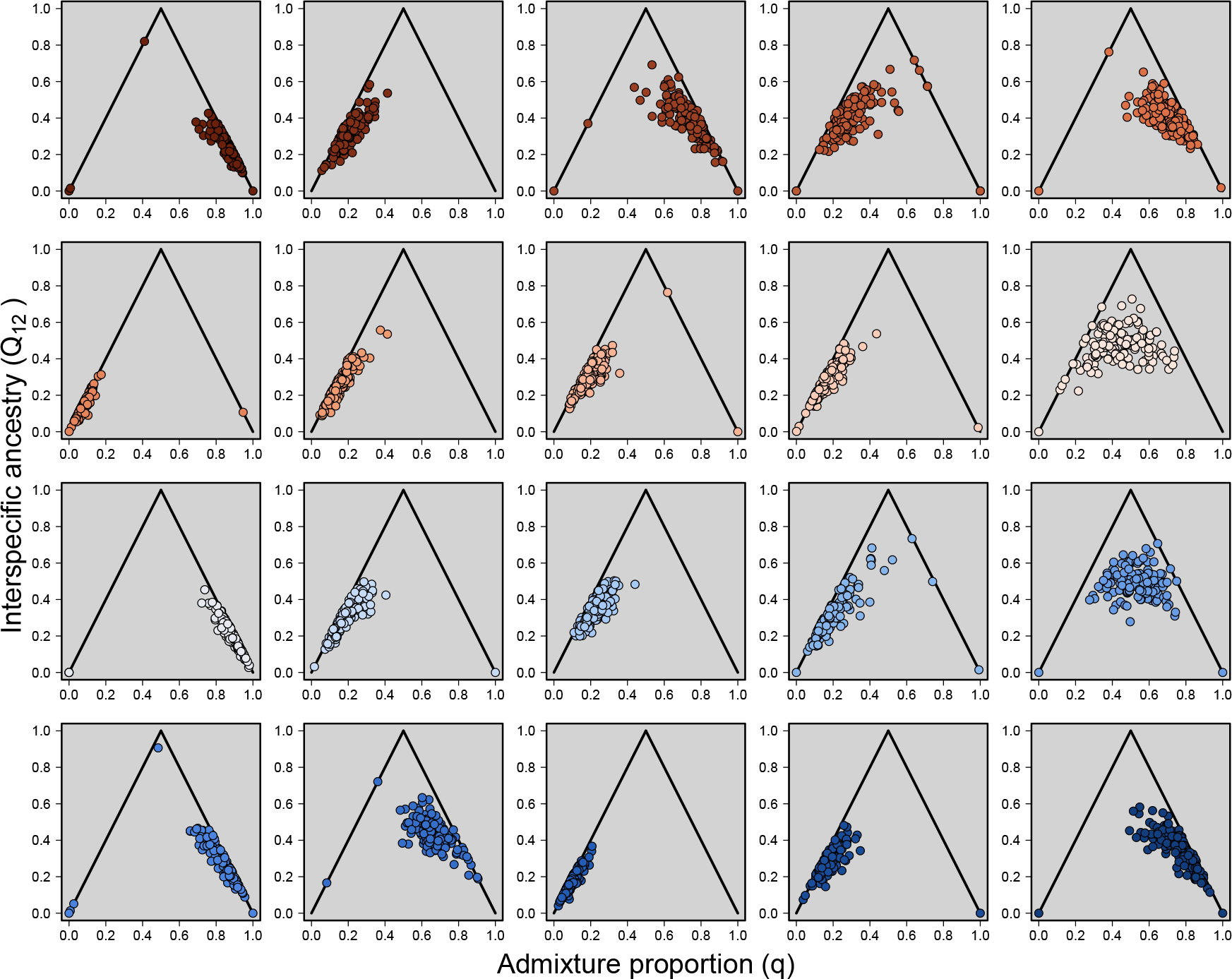
Individual ancestry varies greatly across the twenty replicates from the BDMI, migration (*m*) = 0.01, selection (*c*) = 0.9 simulations (generation 10, deme 6). Paired estimates of admixture proportion (*q*) and inter-specific ancestry (*Q*_12_) can be used to classify individuals as parentals (*q* = 0; *Q*_12_ = 0 or 1), F1 hybrids (*q* = 0.5; *Q*_12_= 1), F2 hybrids (on average *q* = 0.5, *Q*_12_ = 0.5), or backcrosses (residing on the solid black lines) (Gompert *et al*. 2014, Lindtke *et al*. 2014, Shastry *et al*. 2021). Colors correspond to different replicates and match those in other figures.

As many empirical hybrid zones have uneven contributions from each parental population, we explored how much the ratio of the parental populations affected the degree of variation among replicates. Using a three deme structure, we found that replicate explained substantial variation in *q*, *Q*_12_ and the number of junctions, across all ratios of parental populations, after both 10 and 100 generations (Table 1, Fig. S9, Fig. S10, Fig. S11, Fig. S12). There was substantial variation in the F statistics, p values and *R*^2^ estimates (Table 1). However, it does appear, as with the 1:1 parental simulations, that there is a larger effect of replicate when there are BDMIs under strong selection and with low migration between populations (Tables 1, S2 – S6; Figs. S3 – S12).

Each of the three focal loci (1.4: locus under selection; 1.10: locus physically linked to causal locus; 3.4: physically unlinked locus) had different proportions of variation explained by replicate, depending on the simulation scenario. Replicate was a significant predictor of variation in all scenarios for each of the 3 loci (Table S7). The ratio of variance explained by replicate in locus 1.4 ranged from F = 2.7 to 233, where there was little variation explained due to replicate in the BDMI scenario with high migration and moderate selection (*m* = 0.2, *c* = 0.2, F = 2.7, P *<* 0.001, *R*^2^ = 0.017). In contrast, in the BDMI scenario with low migration and high selection, there was huge proportion of variation in genotype explained by replicates (F = 233.1, P *<* 0.001, *R*^2^ = 0.598). Similar patterns across scenarios were found for the other two loci, suggesting that the patterns extended across the genome. We found that using minor ancestry instead of genotype recovered a similar pattern, where replicate explained substantial variation in nearly all scenarios (Table S8). The PCA based on the genotypes from all 510 loci failed to differentiate the replicates from generation 10, except for the scenarios involving a BDMI with *m* = 0.01 and *c* = 0.9 (Fig. S13). At generation 100, however, PCA was able to distinguish among replicates for all low gene flow (*m* = 0.01) scenarios, but replicates were qualitatively identical for all high gene flow (*m* = 0.2) scenarios (Fig. S14). Finally, there was substantial variation in the multinomial regressions of ancestry each of the three focal loci in both the 11 deme scenario (Figs. 5, S15, S16, S17) and the three deme scenario (Fig. S18). When considering how the ratio between parental populations affected consistency, we found that the changes were similar regardless of locus-specific selection. So while consistency changed depending on the ratio between parental populations, this was not consistency due to selection (Fig 6).

**Figure 5:**
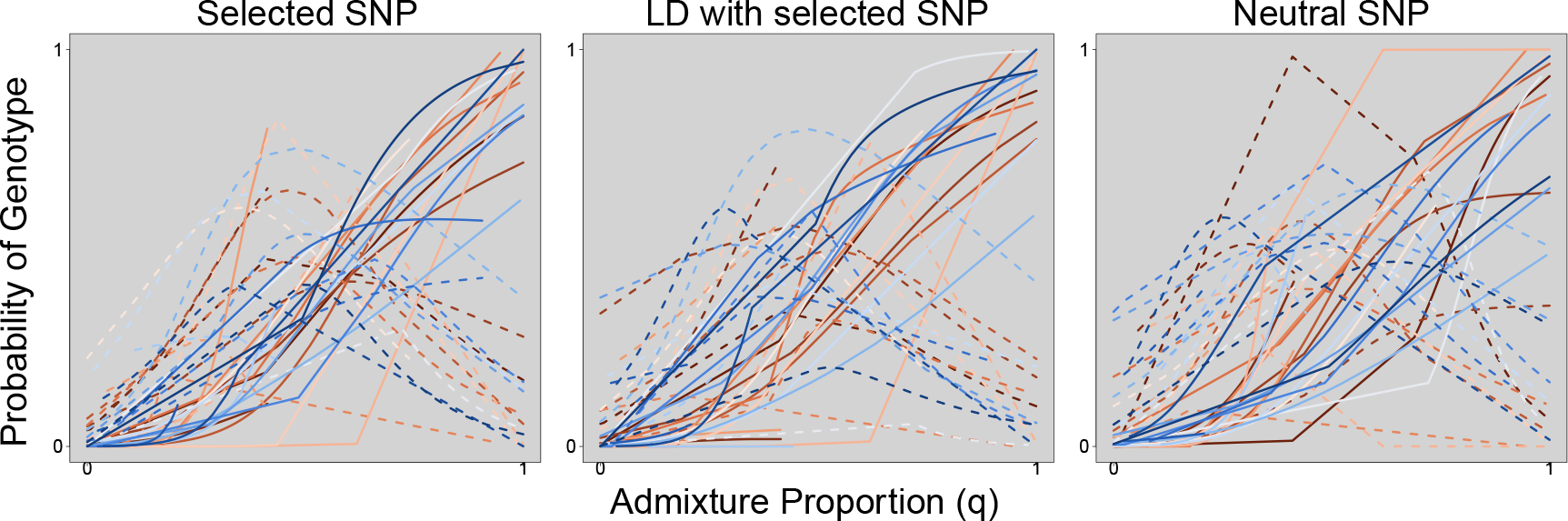
Probability of each AA (homozygous; solid line) and Aa (codominant heterozygote; dashed line) for each of three simulated SNPs in the BDMI scenario where migration is low (*m*=0.01), and selection is high (*c*=0.9) after 10 generations. Colors correspond to different replicates and match those in other figures. This multinomial regression shows the probability of either a homozygous or heterozygous genotype for a given admixture proportion *q*.

**Figure 6:**
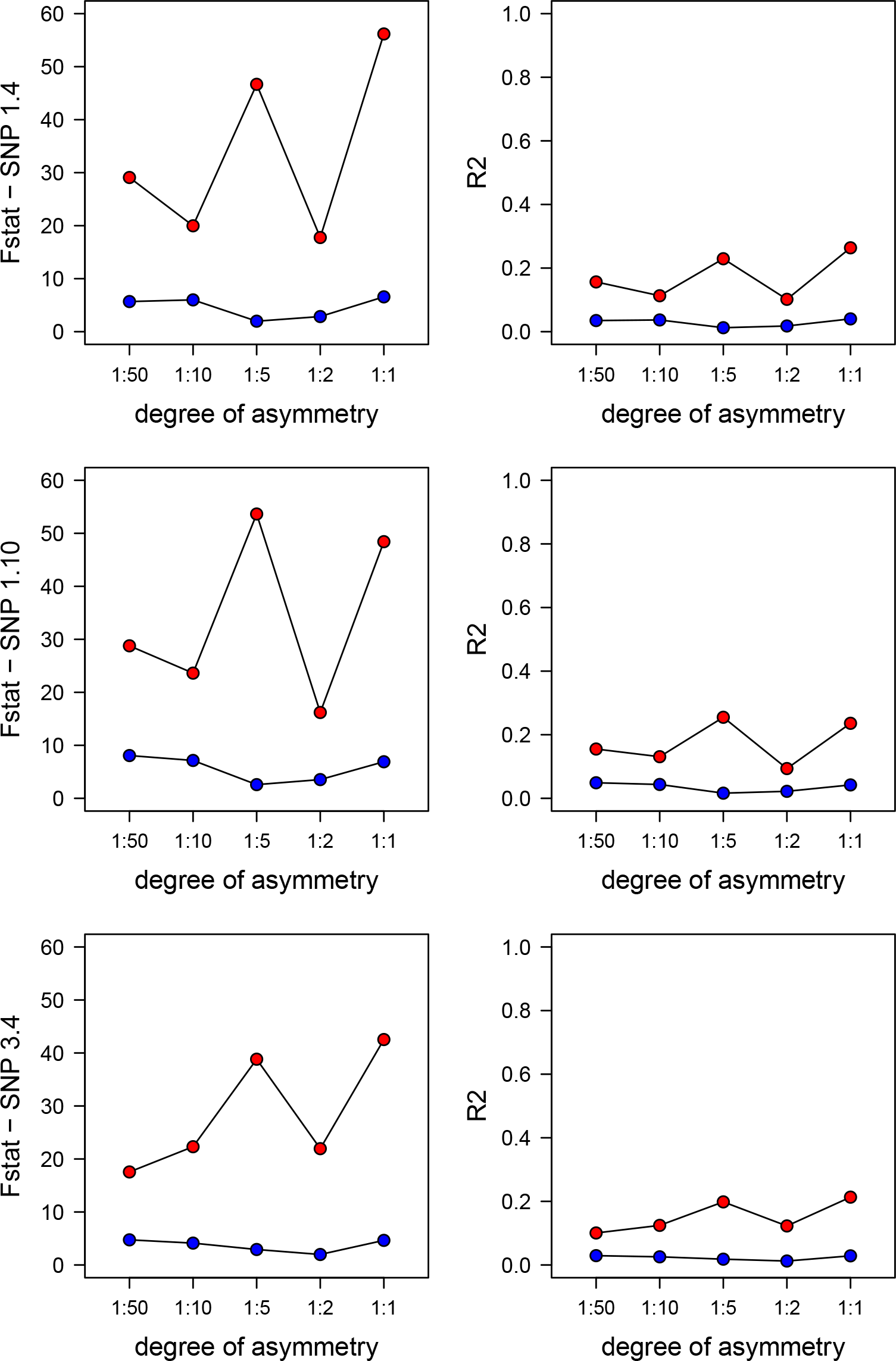
F statistics and *R*^2^ for each locus 1.4 (under selection), locus 1.10 (in linkage with causal incompatibility loci) and locus 3.4 (neutral) from 20 replicates of BDMIs with strong selection (c=0.9), both low and high gene flow across demes (m=0.01 in red and m=0.2 in blue). We varied the ratio of asymmetry between the starting parental population sizes from 50:1 to 1:1 (Table 1). We found that asymmetry affects the consistency across replicates, but that the effect of asymmetry is essentially the same across loci, suggesting that consistency is not indicative of locus specific selection, even when consistency is relatively high.

## Discussion

For many of the evolutionary scenarios we modeled, there was remarkable variation in the outcomes of hybridization across replicates (Fig. 2) despite their identical evolutionary and demographic conditions. Notably, both genome-wide ancestry composition of individuals and genotypes at specific loci varied extensively across replicates. Strong selection on the same loci led to increased genotypic variation among replicates, in contrast to näıve expectations that selection would lead to more consistent outcomes. Moreover, this result extended across the entire genome with pronounced intra- and inter-chromosomal linkage disequilibrium (Fig. 3). Even loci that were not on the same chromosome as the causal loci for incompatibilities were affected by the strong viability selection. For scenarios with unequal starting parental ratios, although outcomes of hybridization were overall more consistent, under strong selection there was still substantial variation among replicates (Fig. 2; Table 1). This lack of a direct causal link between selection strength and outcomes of hybridization suggests caution when tying patterns of consistency to evolutionary processes in replicate hybrid zones.

Similar outcomes of hybridization may occur in the presence or absence of strong selection. We found the least variability explained by among replicate differences when selection was weak and migration was high, suggesting that consistency among replicates could in some cases indicate a lack of reproductive isolation, although other mechanisms could cause consistency too. This was the case regardless of the ratio between parental populations (Table 1), or if minor ancestry was used in analyses instead of genotypes (Table S8). In other words, when hybridization is relatively rare, because migration between populations is low (m=0.01), there is an opportunity for substantial stochasticty leading to variation among replicates. In contrast, when hybridization is common, and selection is not very strong against recombinant individuals, replicate populations are more similar. While it is interesting to note when we might expect replicate hybrid zones to be highly comparable, those populations with little selection against hybrids and lots of hybridization are not terribly relevant from the perspective of identifying loci associated with reproductive isolation, because it is clear that selection on inter-genomic and intra-genomic incompatibilities is not driving hybridization outcomes in these populations.

Variability in outcomes could arise from many causes. Our results highlight a prominent role for stochasticity in the composition of hybrid zones (Blount *et al*. 2008), particularly when selection is strong, meaning that chance events in early generations of hybridization lead to evolution proceeding in dramatically different directions (Gould 1989). Below, we discuss these results in the context of both speciation research and conservation genetics.

### Replicate hybrid zones in speciation genetics

When the geographic ranges of species collide in secondary contact, outcomes of hybridization have been expected to be similar when mechanisms of reproductive isolation are the same. Further, previous work has shown that selection on individual loci can lead these loci to deviate in ancestry from neutral loci elsewhere in the genome (Gompert & Buerkle 2011), though attributing deviation at exceptional loci to drift or selection is fraught (Gompert & Buerkle 2011, McFarlane *et al*. 2021), as many processes can produce this pattern and both drift and selection can contribute to shaping outcomes at once. These assumptions underlie the basic premise of replicate hybrid zone research: similarities across replicates should be indicative of shared isolating barriers, independent of the effects of genetic drift at each individual population, thereby giving insight into the nature of reproductive isolation between a pair of species (Langdon *et al*. 2022, Matute *et al*. 2020, Rieseberg *et al*. 1999, Simon *et al*. 2020, 2021, Westram *et al*. 2021). Despite this expectation, many of our simulations highlight remarkable variation in individual-level parameters (Figs. 2, 4), genotypes (Fig. 3), and introgression (Fig. 5) across replicates in many cases. For example, more than 80% of the variation in *q* is explained by among replicate variation when migration is rare and selection against BDMIs is strong (Table S1). When reproductive isolation is strong, one might anticipate consistency, where strong selection might shore up reproductive isolation. But if there are variable outcomes, then reproductive isolation could collapse in some places but not others, under identical conditions. These unstable equilibria resulting from frequency dependent selection and underdominance (in the context of multi-locus epistasis) in our simulations suggest that a substantial amount of stochasticity could result when reproductive isolation is tested in secondary contact.

In contrast to the idea that strong selection could lead to consistency across replicates, we found that BDMIs combined with strong selection lead to remarkable variation among replicates. Since underdominance created by BDMIs can lead to alternative homozygous genotypic states with full fitness, alternative alleles can rise to high frequency in our different simulation replicates (Fig. 1). This was reflected by a bimodal distribution of *q* scores among replicates particularly when selection on BDMIs was high (Fig. 2). We found that the stronger the barriers to gene flow at specific loci, the more variable were the outcomes under identical conditions, and the more variation was explained by among replicate differences, and thus, the more unstable the allele frequencies are across replicates. Further, any drift will lead to alternative homozygotes being favoured due to frequency dependent selection (Ayala & Campbell 1974, Heino *et al*. 1998, Gompert *et al*. 2012). This means that small perturbations to identical systems due to stochasticity can result in drastically different outcomes (Fig. 1). The unstable equilibrium dynamics predominate in the first few generations after secondary contact, but eventually steady migration will bring hybrid zones back to migration-drift balance. In the meantime, selection for either parental homozygote has set the trajectory of the system for later generations. This is why late generations were well predicted by early generations (Figs. S7, S8). This is consistent with previous work that has demonstrated the sensitivity of evolutionary trajectories to starting conditions (Simões *et al*. 2008, Losos 2011) and the historical sequence of events (i.e. historicity; Lewontin 1966); while the replicates in our simulations have identical conditions, stochasticity and drift lead to differences in early generations that are perpetuated.

We expect to see the least stochasticity when comparing replicates from scenarios with unequal starting numbers of individuals from admixed populations, as the outcomes of hybridization (i.e. which pair of alleles go to fixation) should be more easily be predicted based the major allele frequency (Kimura 1962). Thus, there could be increased consistency, but it would not be because of selection. Indeed, this is what we found (Fig. 2). We tested how changing the starting ratios of the parental populations affected consistency among replicates, and we found that there was generally more consistency when there was more asymmetry (Fig. 6). However, we have also shown that this increase in consistency at particular loci was when selection was low, and was likely due to drift. If selection led to more consistency across replicates than drift did, which would allow consistency to be used to infer selection, this would be most likely be seen in the 1:1 scenarios. In empirical studies with asymmetry, we expect that observed consistency would likely be due to drift, not selection, and that in these cases it would be even more difficult to point to selection as the underlying mechanism.

Our conception of speciation typically includes a set of phenotypes and underlying genotypes that contribute to reproductive isolation between diverged taxa. However, the results of our simulation study suggest that if taxa hybridize and make fertile and viable progeny in multiple instances of secondary contact, a single, consistent set of loci contributing to reproductive isolation would lead to variable evolutionary dynamics of hybrids and gene flow. That variability will make it more challenging to map the genes associated with reproductive isolation, even if there is a single, consistent genetic architecture. Indeed, we have previously noted that genetic drift can lead to patterns in the genome that can mimic the predicted outcomes of selection in secondary contact (Gompert & Buerkle 2011, McFarlane *et al*. 2021). Beyond the challenge of mapping pattern to process, the variation in the effectiveness of reproductive isolation even under (unrealistically) constant conditions suggests that the nature of isolation and gene flow between species is more complex than we have typically appreciated, or than would be suggested by a simple interpretation of ‘speciation genetics’. For these reasons, we suggest that understanding the variation in effectiveness of incompatibilities, where secondary contact occurs in more than one location (or time point), is an interesting avenue for future research. Additionally, coupling between barrier effects, including loci involved in pre- or postzygotic isolation, has been predicted to increase consistency across clines within a hybrid zone (Barton 1983, Butlin & Smadja 2018). While we have not considered coupling directly here, we predict that this strengthening of reproductive isolation via weak coupling could increase variability across replicate hybrid zones, as the potential for stochasticity acting on multiple traits would likely increase the variability, not decrease it. However, when we take a strict interpretation of coupling as only between genetic barriers (Barton 1983), and not when any two potential barriers act at once (Butlin & Smadja 2018), we might expect that coupling would decrease variability between replicates.

### Implications for conservation research

In the realm of conservation, where replicate hybrid zones are studied because of the potential negative consequences of hybridization between endemic and introduced species (Allendorf *et al*. 2001, Mandeville *et al*. 2017, Todesco *et al*. 2016, Wolf *et al*. 2001), our findings are somewhat challenging. Given finite budgets for conservation, the ability to generalize from one setting with potentially hybridizing individuals to other unsampled settings (i.e. out-of-sample prediction) would be extremely useful. However, except in specific circumstances (e.g. when selection against hybrids is known to be low, and hybridization is common), we anticipate substantial variation among hybrid zones even under identical demographic and evolutionary processes, if hybridization is driven by genomic incompatibilities. This means that to assess the likely extent of hybridization in each population, for example to protect individuals in those settings with the least hybridization, each locality might need to be sampled. Since this pattern is driven by genetic drift, the variation among localities will be even higher with small effective population sizes, as would be expected if one species has been introduced to the range of another. However, our results varying the starting parental ratios (Fig. 2, Table 1) suggest that when parental species ratios are uneven at the start of hybridization, results are relatively consistent in the absence of strong selection. Note, however, that these simulations explicitly exclude ecological factors, which could also produce variable hybridization outcomes.

The predictability of evolution is negatively (but not linearly) related to effective population size (Szendro *et al*. 2013), so when considering small, introduced or endangered populations, practitioners must assume that drift and the corresponding stochasticity in the composition of populations are even stronger. One optimistic point worth highlighting is the consistency found across time within replicate hybrid zones. If conditions do not change over time, the distribution of admixture proportions in generation 100 is highly correlated with the distribution in generation 10 (Figs. S7, S8). Practitioners will need to weigh the need for temporal replication to ‘check in’ relative to the need for spatial replication, based on our finding that every new hybrid zone could have a different distribution than those already sampled. However, one caveat is that this temporal stability is based on a system with constant conditions, which is fairly unrealistic.

### Caveats

One difference between our study and some empirical studies is the use of minor ancestry at haplotypes, rather than admixture proportion, to determine to what extent there are regions of the genome with depleted variation, suggesting incompatibilities (e.g. Langdon *et al*. 2022, Schumer *et al*. 2018). While we explored this for specific loci (Table S8), we are conscious that polarizing by minor parent assumes that backcrossing in either direction has the same consequences, regardless of the ancestry of the genetic background. Given that most of our simulations assumed the same parental population size and equal contributions to admixture (i.e. there was no asymmetry), the population determined to be the minor ancestor varied among replicates (Fig. 1). When we compared minor ancestry at specific loci, we found a very low correlation between (arbitrarily chosen) replicate 1 and 2 and significant variation across replicates in nearly all scenarios (Table S8), suggesting that using minor ancestry rather than genotype does not substantially decrease the variation between replicates.

A second key difference between our simulations and the expectations in empirical systems is the heterogeneous recombination landscape that has been demonstrated to affect speciation (Nachman & Payseur 2012). Our simulations have a homogenous recombination landscape based on a Poisson process (Lindtke & Buerkle 2015). This homogenous recombination means that recombination hotspots and cool spots, which could lead to purging of even beneficial alleles in empirical systems (Moran *et al*. 2021), have not been simulated here. This also means that we would not have found a correlation between recombination and minor ancestry (as in Langdon *et al*. 2022), although we also did not find a correlation of minor ancestry between replicate populations. Future work could simulate such recombination landscapes and ask how this affects variation in outcomes across replicate hybrid zones.

A third caveat is that we have not simulated adaptive introgression, where an allele from one population introgresses through the hybrid zone into the other population due to directional selection. We expect that some of the same problems we have demonstrated will apply in that circumstance as well, even if underdominance does not drive variation as we expect it does for BDMIs. Some empirical studies have found evidence of adaptive introgression from introduced species, using replicate hybrid zones (e.g., Bay *et al*. 2019, Calfee *et al*. 2020), but few have explicitly ruled out genetic drift as a mechanism, and selection is regularly implicated even when hybrid zones do not show concordant patterns (but see Nouhaud *et al*. 2022). Given that adaptive introgression is only expected to be possible when recombination is quite high, allowing a beneficial allele to be separated from a neutral or deleterious haplotype (Moran *et al*. 2021, Uecker *et al*. 2015), the *a priori* expectation must be that introgression is commonly due to non-adaptive forces (*sensu* Barrett & Hoekstra 2011).

Finally, more generally, our results are illustrative of variability that arises in a particular set of circumstances and are not meant to be a perfect reflection of the most likely population structure and fitnesses in secondary contact for a pair of taxa in nature. While we have perturbed these simulations to examine robustness, such as considering both an 11 deme and 3 deme population structure (Lindtke & Buerkle 2015; Figs. S2, S3), simulations can never cover all possible biological variability. The expectations for any natural setting should be informed by knowledge of the system, which will include an appreciation of the complexities due to chance in finite populations, unstable equilibrium allele frequencies (and ancestries), and the dynamics of gene flow between populations with potentially unequal and variable size through time.

### Strategies for analyses of replicate hybrid zones

Even when there is substantial variability across hybrid zones, there is still the opportunity to capitalize on replication when available. This might be particularly fruitful when the genetic architecture of incompatibilities is expected to be polygenic or selection is less strong, as such cases have less variation explained by among replicate differences (Table S1). Given that polygenic architectures are likely responsible for many cases of reproductive isolation (Barton 2022), we are encouraged by our results. By accounting statistically for this variability across replicates (rather than assuming all replicates are comparable), one can examine both global and local patterns. For example, some studies have used replicate hybrid zones to ask how ecological variation affects distributions of admixture (e.g. Chatfield *et al*. 2010, Culumber *et al*. 2012, Mandeville *et al*. 2019, 2021, Velo-Antón *et al*. 2021), a question that assumes variation across replicates. Here, we have used ANOVA to estimate variance due to replicate, but we have not incorporated other variables (e.g. environmental variability) into our analyses. ANOVAs are excellent for our balanced simulations, but might not be the appropriate method for studies that want to account for variability among replicates and also ask how some other variable (e.g. human activity, temporal variation) affects hybridization outcomes. Some studies of replicate hybrid zones have used PCA or similar approaches to assess to what extent there were substantial differences among replicates, and then using pairwise correlations between studies, to determine to what extent there was concurrence between admixture distributions, allele frequencies, or cline parameters (e.g. Westram *et al*. 2021). We found that PCAs did not differentiate among replicates well in the 10th generation after admixture (Fig. S13), suggesting that PCAs are not sufficient to confirm similarity among replicates.

One could build more sophisticated hierarchical models beyond ANOVA, but the usual rules about sample size for hierarchical models will apply. For example, Calfee *et al*. (2020) fit a classic logistic cline model to inferred genome-wide individual ancestry proportions, which fit a cline specific to two replicate instances of the escape of *Apis mellifera scutellata* and subsequent hybridization with *Apis mellifera*. They were thus able to make inferences about the similarities and differences of these two hybrid zones, while accounting for between replicate variation. By using a hierarchical model, which allows for differences among populations in genome wide distributions, but assumes that these distributions are all pulled from a global population, there is an opportunity to compare clines at the same locus across populations, without assuming that the populations are perfectly comparable. This means that different taxa and systems will be more fruitful for understanding causes of different outcomes of hybridization, such as multiply introduced species like trout (Mandeville *et al*. 2019), mussels (Simon *et al*. 2020), and organisms that are amenable to laboratory settings where there can be substantial replication (e.g. *Saccharomyces* sp., Dunn *et al*. 2013; *Tigriopus californicus*, Pritchard & Edmands 2013). There are opportunities to use replicate hybrid zones to shed light on both local and global patterns of hybridization and introgression, while being cautious to not assume that replicates will be predictive of other replicates or unsampled sites.

One challenge when studying hybridization across spatial replicates is to think about how independent replicate sites are. At one end of the spectrum, ‘replicates’ may not be independent at all for study systems that have high levels of gene flow over the spatial scale of sampling. In contrast, some hybrid zones may be truly independent, such as instances of secondary contact isolated from one another in a mosaic hybrid zone or by a substantial distance. For example, bighorn sheep have been reintroduced to several isolated mountain ranges across North America using founders from multiple source ancestries (Jahner *et al*. 2019). More often, studies may involve multiple transects of secondary contact between diverged lineages, as in studies of house mice in Europe (Bímová *et al*. 2011, Payseur *et al*. 2004, Teeter *et al*. 2010) or *Gryllus* crickets in North America (Larson *et al*. 2014). Replicated hybrid zones may also have complex spatial structure, and replicates may be hierarchically nested, as is the case with replicating across fish hybrid zones in small streams within a larger river basin, and then also replicating across basins (e.g. *Catostomus* sucker hybridization or rainbow *×* cutthroat trout hybridization; Mandeville *et al*. 2017, Muhlfeld *et al*. 2017). Geographic distribution of hybridizing species, as well as the dispersal mode and capacity of study organisms, are likely to be quite important in defining independence of replicate sites, and may influence decisions about statistical treatment of comparisons. We emphasize that studies of replicate hybrid zones at all levels of independence are likely to be informative, but clearly defining the independence of replicates is an important goal.

## Conclusion

This research was motivated by our own ambitions to use replicate hybrid zones to ask basic and applied questions in our study systems. Indeed, we have written about the promise of replicate hybrid zones in our own papers (Buerkle & Lexer 2008, Jahner *et al*. 2021, Lindtke *et al*. 2012, Mandeville *et al*. 2017, McFarlane *et al*. 2021), so we were fairly confident that we could link the outcomes of hybridization to the evolutionary processes that generated them, at least for some scenarios. As it turns out, early generation stochasticity can dramatically alter the outcomes of hybridization across hybrid zones, leading to extreme variability across replicates with identical starting conditions. Moreover, the degree of variability is amplified with increasing strength of selection on incompatibilities, suggesting that the ‘simplest scenarios’ will be the most difficult in which to tie pattern to process. Despite these results, we believe the future of replicate hybrid zone research remains promising, particularly if potential reasons for variation among hybrid zones are considered appropriately. We hope this study will encourage 1) the development of new statistical models that explicitly account for variability across hybrid zones and use this variability to better understand evolutionary and demographic causes of reproductive isolation; 2) future modeling exercises that examine the countless other important axes of variability that were not explored here; and 3) new empirical studies of replicate hybrid zones that acknowledge the potential lack of identifiability of evolutionary processes based on our observed outcomes of secondary contact.

## Acknowledgements

We are grateful to Cassandre Pyne, Amanda Meuser, Amy Pitura, Teaghan Frauley, Chris Nice, and Zach Gompert for previous discussions of this manuscript. We are also thankful for the comments of four thoughtful reviewers. This research was supported by the Modelscape Consortium with funding from NSF (OIA2019528). Analyses were supported by the University of Wyoming’s Advanced Research Computing Center and its Beartooth Computing Environment, Intel x86 64 cluster (https://doi.org/10.15786/M2FY47).

## Data Accessibility Statement

The original source code for the simulation software dfuse can be found at the Dryad Digital Repository associated with Lindtke & Buerkle (2015) (doi: 10.5061/dryad.0506g). All code and input files for analyses can be found at on Zenodo https://zenodo.org/account/settings/github/repository/erynmcfarlane/predict_hybrids, (McFarlane *et al*. 2022).

## Benefit-Sharing Statement

Benefits from this research accrue from the sharing of our data and results on public databases as described above.

## Author contributions

SEM and EGM had the initial idea to think carefully about the utility of replicate hybrid zones in evolution and conservation. DL and SEM ran all of the simulations. SEM, JPJ, and CAB went down a deep, statistical rabbit hole before ultimately deciding to use ANOVA to analyze everything. SEM wrote the initial draft of the manuscript, and all authors contributed to subsequent revisions.

## Notes

### Competing Interest Statement

The authors have declared no competing interest.

### Summary of Updates

This version fully incorporates simulations that vary the ratio between parental populations to hybrid zones in a variety of scenarios. This was done in response to reviewer comments.

