## Supplementary Materials for "Selection leads to remarkable variability in the outcomes of hybridization across replicate hybrid zones"

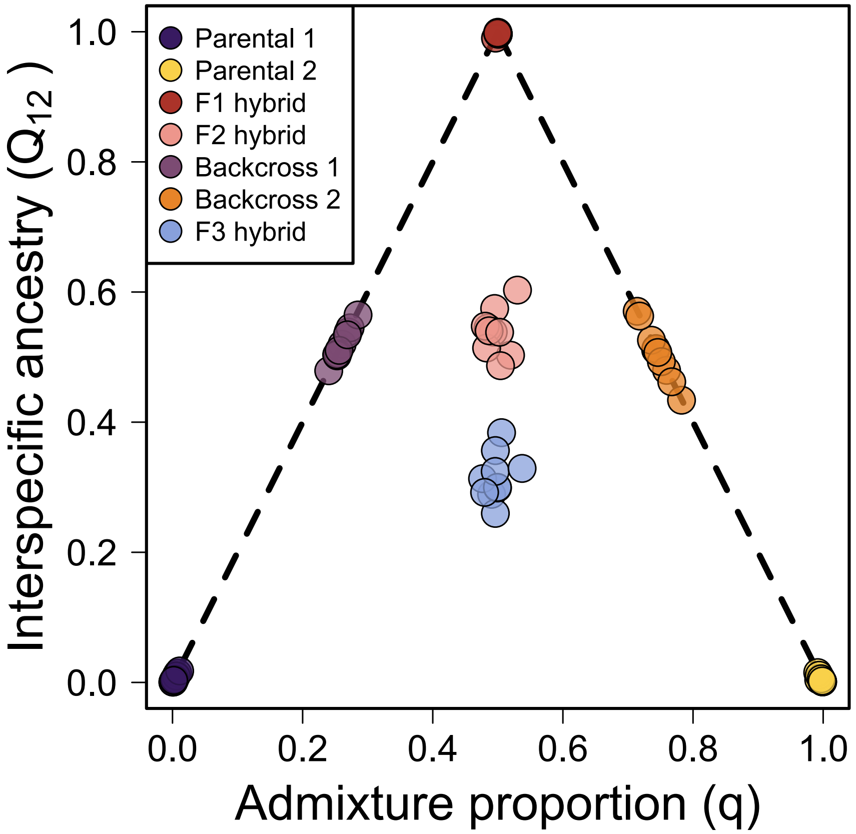

Figure S1: An individual's hybrid status can be classified based on the distribution of admixture proportion ($q$) and interspecific ancestry ($Q_{12}$). Figure modified from Shastry et al. (2021).

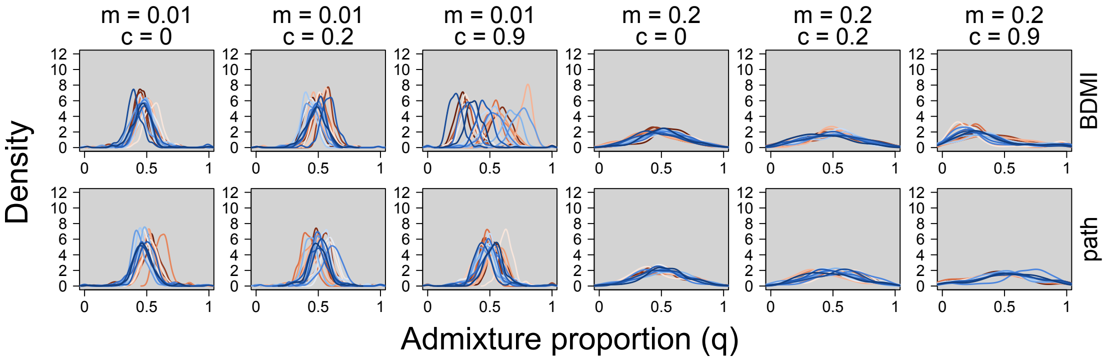

Figure S2: Selection leads to remarkable variation in the outcomes of hybridization across replicate hybrid zones. The distribution of inter-specific ancestry ($q$) from generation 10 is shown for 20 replicates from the 12 scenarios that varied by migration ($m$), selection ($c$) and incompatibility (BDMI versus pathway), in the three deme system, as contrasted with the 11 deme system presented in the main manuscript. Colors correspond to different replicates and match those in other figures.

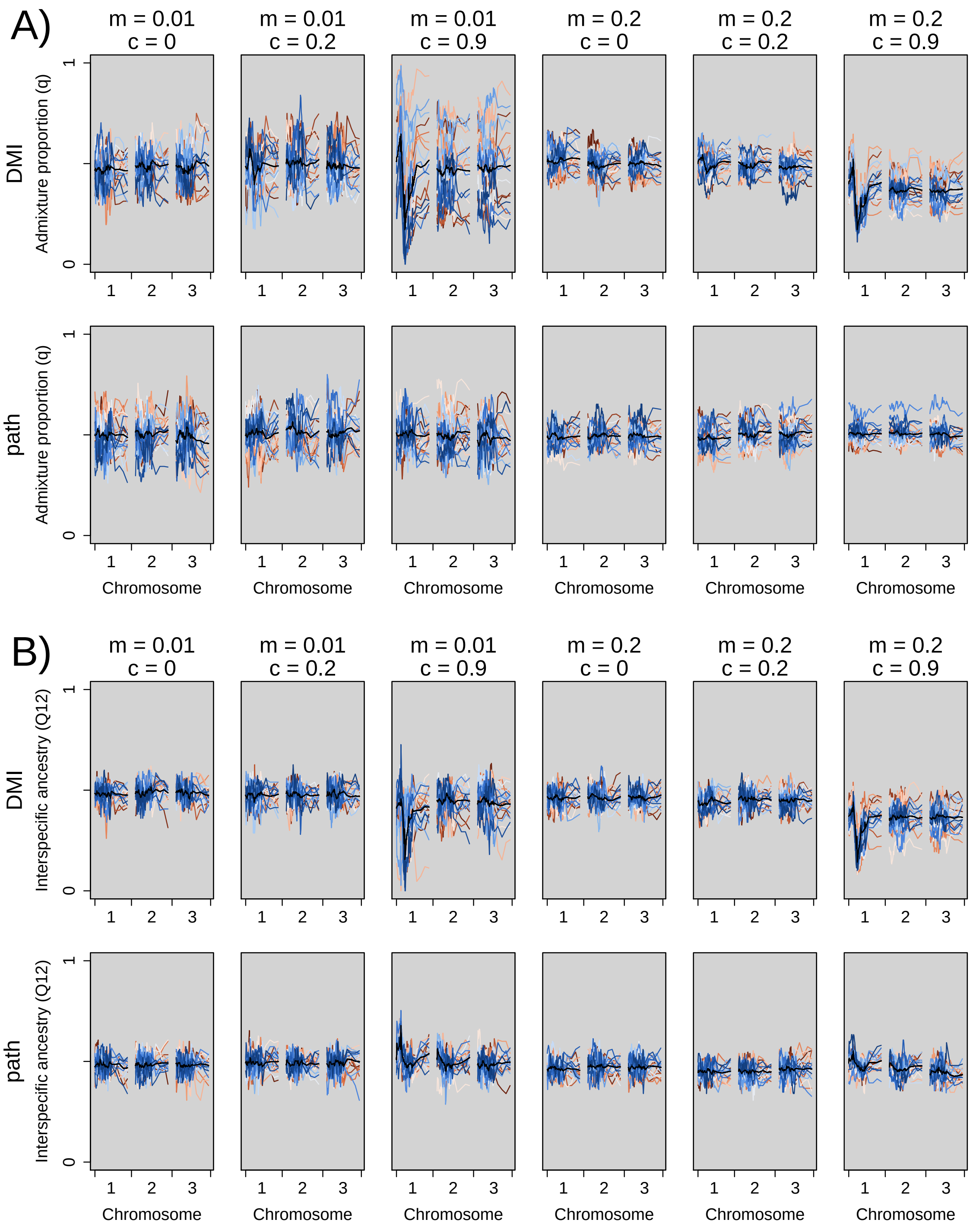

Figure S3: Locus-specific variation in average (among individuals in a replicate) admixture proportion ($q$, panel A) and inter-specific ancestry ($Q_{12}$, panel B) for replicates (different colours), six combinations (columns) of migration ($m = 0.01$ or $0.2$) and selection ($c = 0$, $0.2$ or $0.9$), and two types of incompatibilities (BDMI or pathways, in rows), as simulated in a 3 deme system. Colors correspond to different replicates and match those in other figures. The black line is the average across all replicates, as in Lindtke and Buerkle 2015. The plots are for simulations after ten generations. Chromosome 1 has loci with BDMI or pathway incompatibilities, chromosome 2 has pathway incompatibility loci, while chromosome 3 has only neutral loci, and is representative of the rest of the genome (chromosomes 4–10).

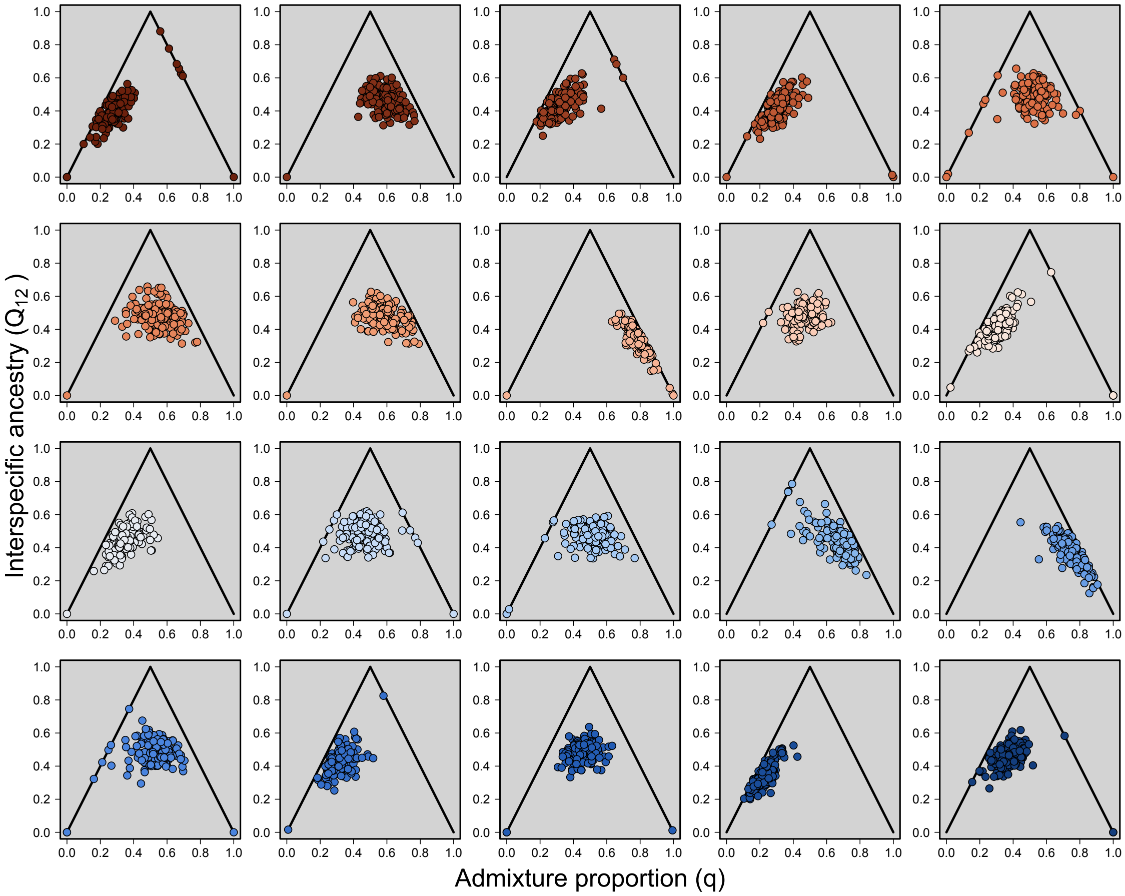

Figure S4: Individual ancestry varies greatly across the twenty replicates from the BDMI, migration ($m$) = 0.01, selection ($c$) = 0.9 simulations (generation 10, deme 6) in a 3 deme system. Paired estimates of admixture proportion ($q$) and inter-specific ancestry ($Q_{12}$) can be used to classify individuals as parentals ($q$ = 0; $Q_{12}$ = 0 or 1), F1 hybrids ($q$ = 0.5; $Q_{12}$= 1), F2 hybrids (on average $q$ = 0.5, $Q_{12}$ = 0.5), or backcrosses (residing on the solid black lines; Gompert et al. 2014, Lindtke et al. 2014, Shastry et al. 2021). Colors correspond to different replicates and match those in other figures.

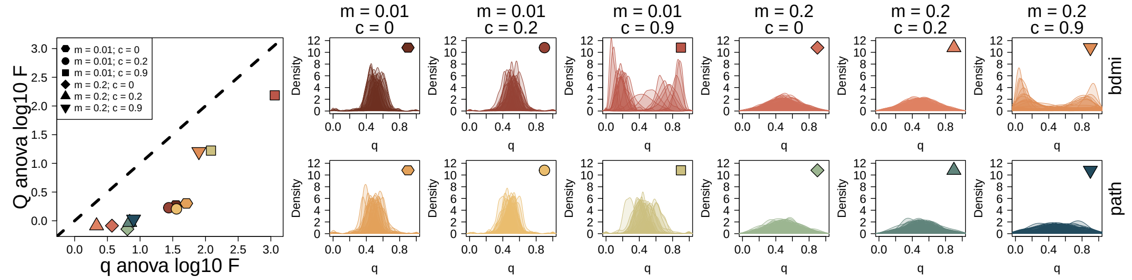

Figure S5: Replicate hybrid zones were typically more variable when selection was high, migration was low, and the genetic architecture was characterized by a BDMI. Anova was used to assess whether freplicate simulations differed from one another for a given scenario [migration ($m$); selection ($c$); architecture (BDMI versus pathway); environment (env)]. The large left panel displays ANOVA F statistics from models including admixture proportion ($q$) and ancestry complement ($Q$), while the smaller panels on the right show the distribution of $q$ for each of the twenty replicates for each scenario (generation 10, deme 6).

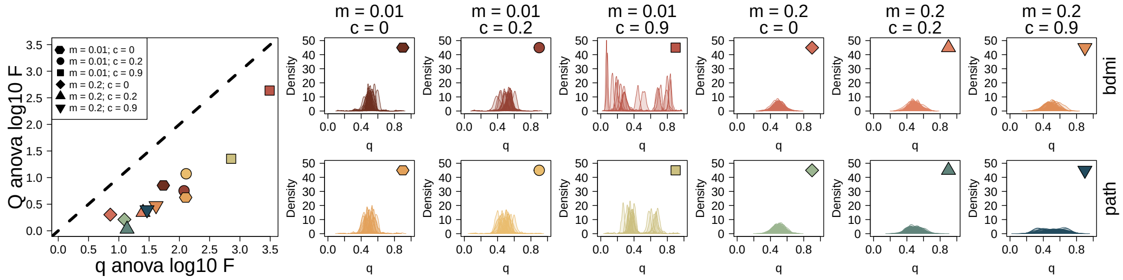

Figure S6: Replicate hybrid zones were typically more variable when selection was high, migration was low, and the genetic architecture was characterized by a BDMI. Anova was used to assess whether replicate simulations differed from one another for a given scenario [migration ($m$); selection ($c$); architecture (BDMI versus pathway); environment (env)]. The large left panel displays ANOVA F statistics from models including admixture proportion ($q$) and ancestry complement ($Q$), while the smaller panels on the right show the distribution of $q$ for each of the twenty replicates for each scenario (generation 100, deme 6).

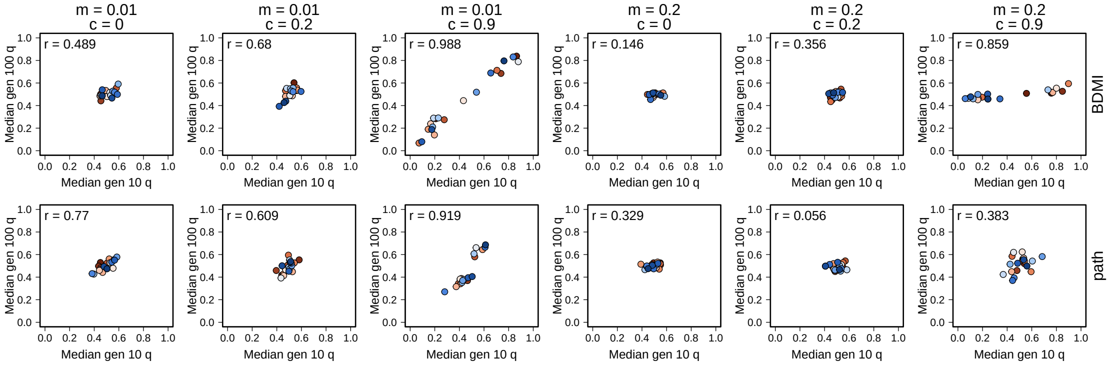

Figure S7: While a replicate's median admixture proportion ($q$) in generation 100 is associated with $q$ in generation 10 (Pearson's correlation; r), the strength of the relationship varies by scenario [migration ($m$); selection ($c$); architecture (BDMI versus pathway); environment (env)].

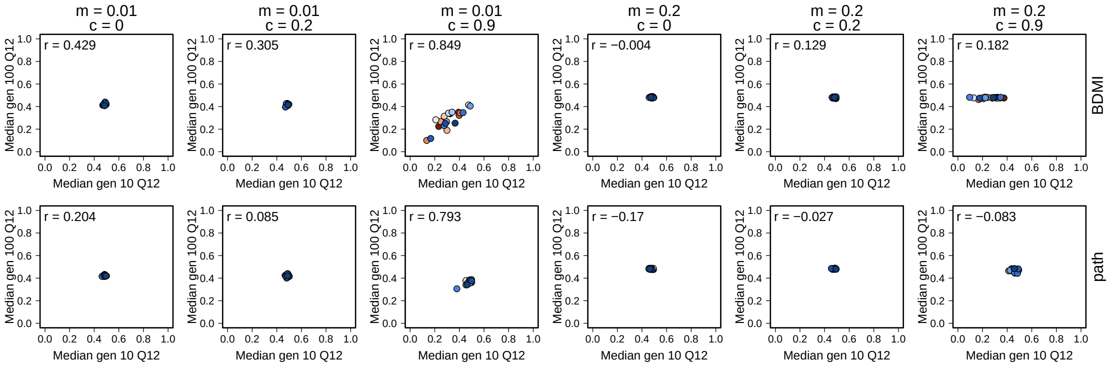

Figure S8: While a replicate's median ancestry complement ($Q_{12}$) in generation 100 is associated with $Q_{12}$ in generation 10 (Pearson's correlation; r), the strength of the relationship varies by scenario [migration ($m$); selection ($c$); architecture (BDMI versus pathway); environment (env)].

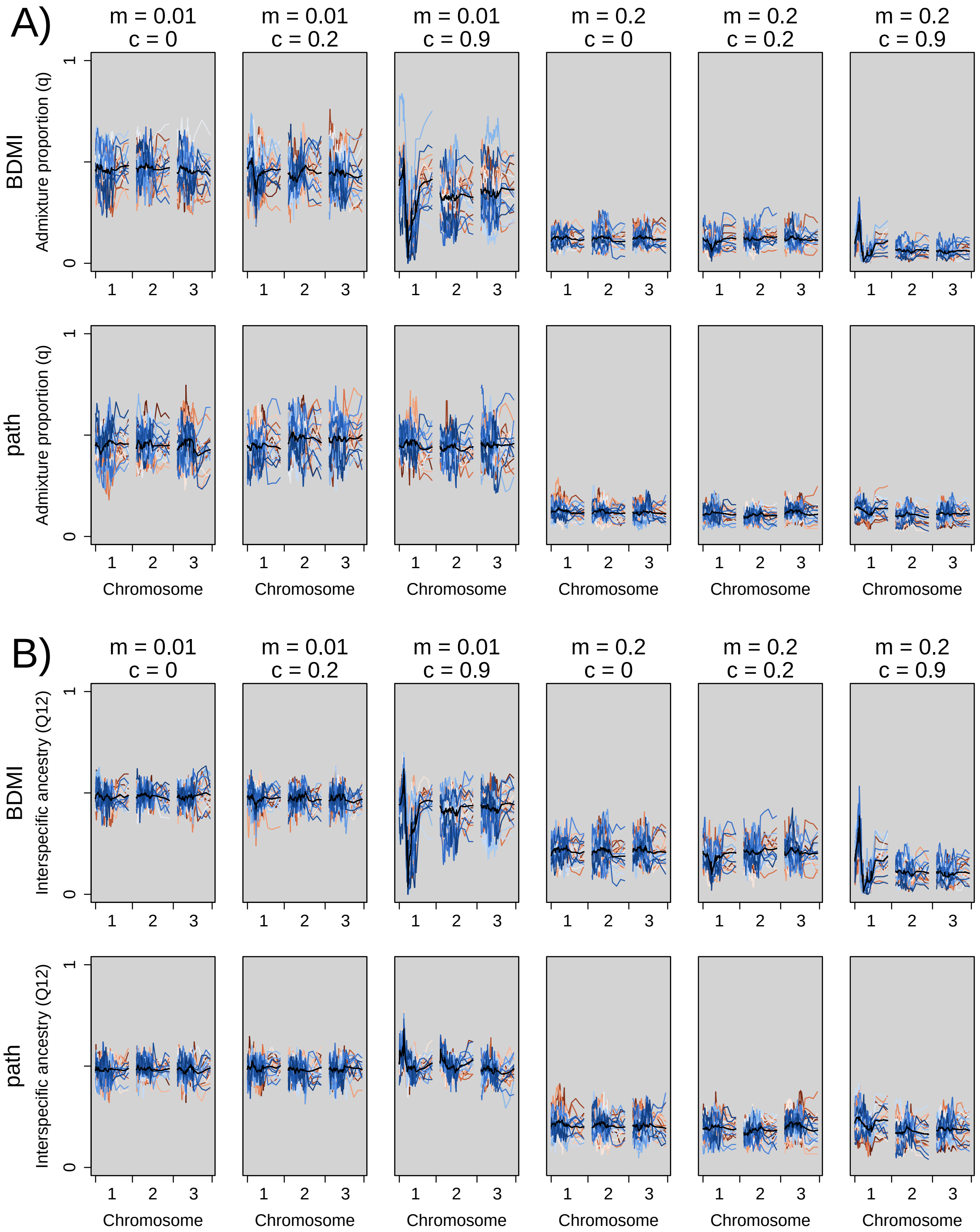

Figure S9: This figure shows results for simulations with parental species at initial ratios of 10:1, in contrast to Fig. 3 in the main text, which shows parental ratios of 1:1. Locus-specific variation in average (among individuals in a replicate) admixture proportion ($q$, panel A) and inter-specific ancestry ($Q_{12}$, panel B) for replicates (different colours), six combinations (columns) of migration ($m = 0.01$ or $0.2$) and selection ($c = 0$, $0.2$ or $0.9$), and two types of incompatibilities (BDMI or pathways, in rows), as simulated in a 3 deme system. Colors correspond to different replicates and match those in other figures. The black line is the average across all replicates, as in Lindtke and Buerkle 2015. The plots are for simulations after ten generations. Chromosome 1 has loci with BDMI or pathway incompatibilities, chromosome 2 has pathway incompatibility loci, while chromosome 3 has only neutral loci, and is representative of the rest of the genome (chromosomes 4–10).

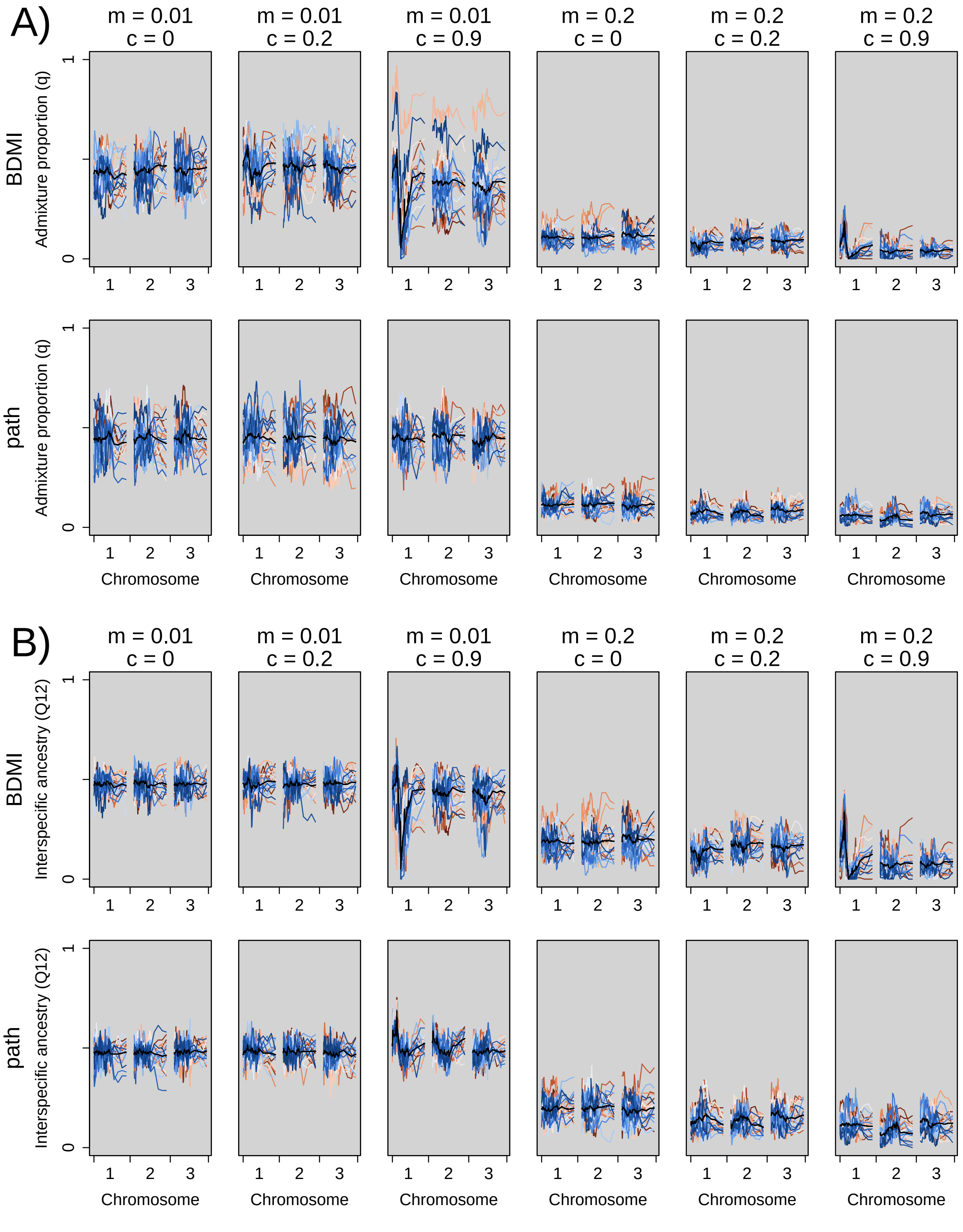

Figure S10: This figure shows results for simulations with parental species at initial ratios of 50:1, in contrast to Fig. 3 in the main text, which shows parental ratios of 1:1. Locus-specific variation in average (among individuals in a replicate) admixture proportion ($q$, panel A) and inter-specific ancestry ($Q_{12}$, panel B) for replicates (different colours), six combinations (columns) of migration ($m = 0.01$ or $0.2$) and selection ($c = 0$, $0.2$ or $0.9$), and two types of incompatibilities (BDMI or pathways, in rows), as simulated in a 3 deme system. Colors correspond to different replicates and match those in other figures. The black line is the average across all replicates, as in Lindtke and Buerkle 2015. The plots are for simulations after ten generations. Chromosome 1 has loci with BDMI or pathway incompatibilities, chromosome 2 has pathway incompatibility loci, while chromosome 3 has only neutral loci, and is representative of the rest of the genome (chromosomes 4–10).

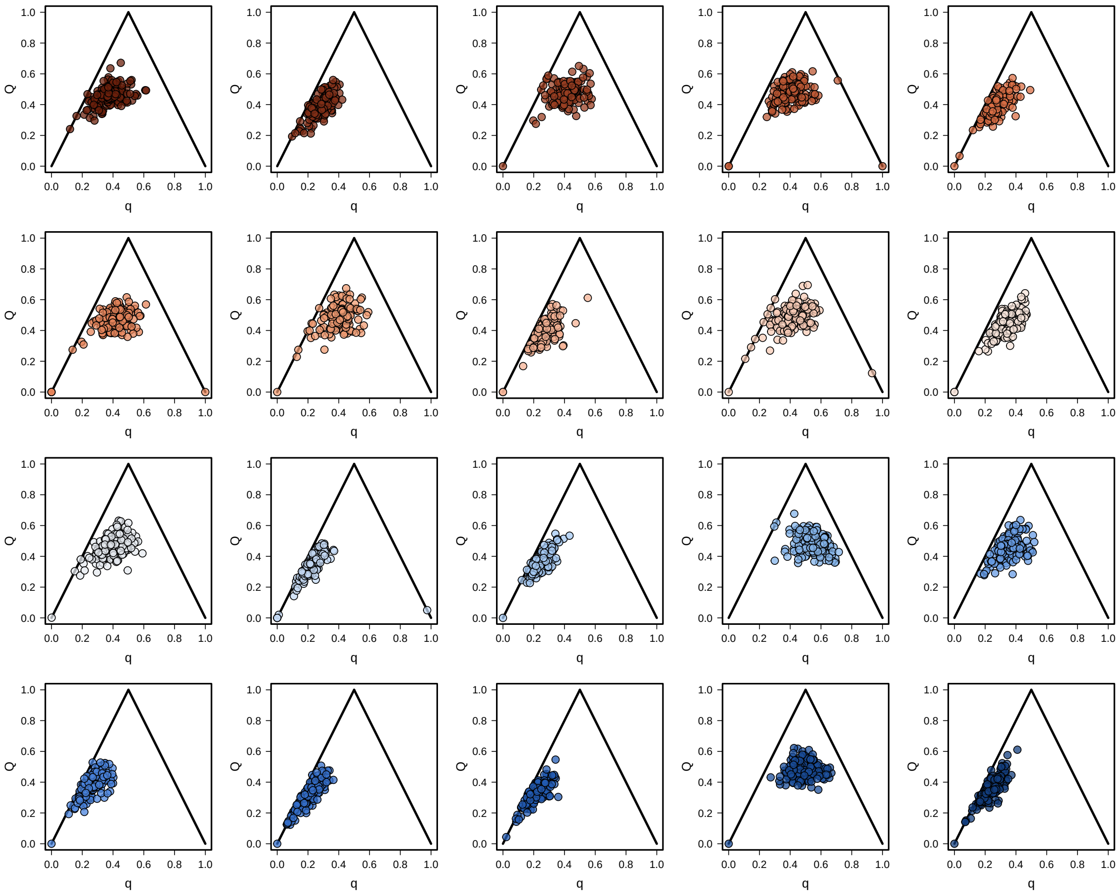

Figure S11: This figure shows simulations with parental species at initial ratios of 10:1, in contrast to main text Fig. 4, which shows parental species at a 1:1 ratio. Individual ancestry varies greatly across the twenty replicates from the BDMI, migration ($m$) = 0.01, selection ($c$) = 0.9 simulations (generation 10, deme 6) in a 3 deme system. Paired estimates of admixture proportion ($q$) and inter-specific ancestry ($Q_{12}$) can be used to classify individuals as parentals ($q$ = 0; $Q_{12}$ = 0 or 1), F1 hybrids ($q$ = 0.5; $Q_{12}$= 1), F2 hybrids (on average $q$ = 0.5, $Q_{12}$ = 0.5), or backcrosses (residing on the solid black lines; Gompert et al. 2014, Lindtke et al. 2014, Shastry et al. 2021). Colors correspond to different replicates and match those in other figures.

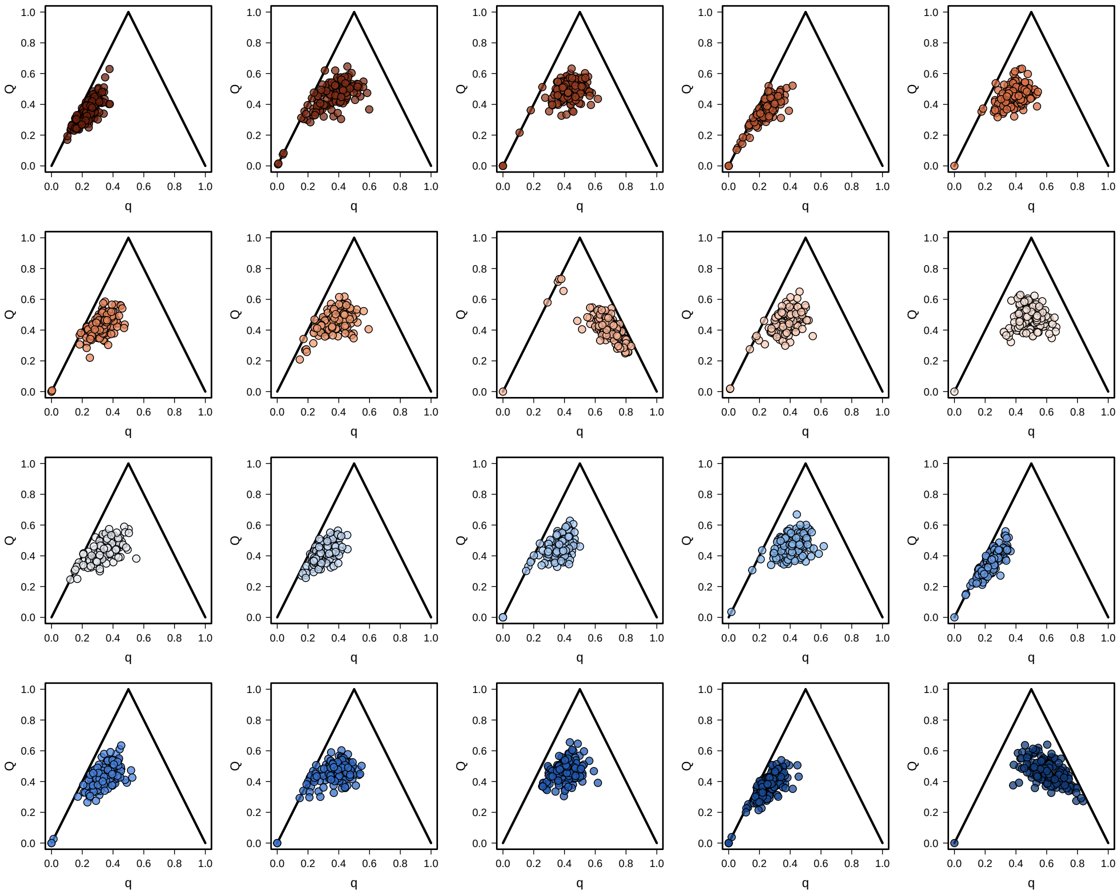

Figure S12: This figure shows simulations with parental species at initial ratios of 50:1, in contrast to main text Fig. 4, which shows parental species at a 1:1 ratio. Individual ancestry varies greatly across the twenty replicates from the BDMI, migration $(m) = 0.01$, selection $(c) = 0.9$ simulations (generation 10, deme 6) in a 3 deme system. Paired estimates of admixture proportion $(q)$ and inter-specific ancestry $(Q_{12})$ can be used to classify individuals as parentals ($q = 0$; $Q_{12} = 0$ or 1), F1 hybrids ($q = 0.5$; $Q_{12}= 1$), F2 hybrids (on average $q = 0.5$, $Q_{12} = 0.5$), or backcrosses (residing on the solid black lines; Gompert et al. 2014, Lindtke et al. 2014, Shastry et al. 2021). Colors correspond to different replicates and match those in other figures.

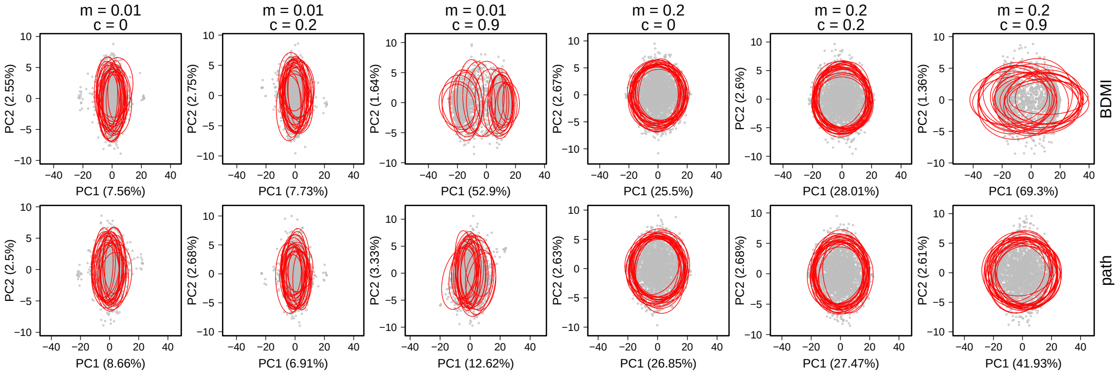

Figure S13: Principal component analysis (PCA) on individual genotypes was used to assess whether replicate simulations differed from one another for a given scenario [migration ($m$); selection ($c$); architecture (BDMI versus pathway); environment (env)]. Gray points correspond to individual scores for the first two PCs, and red ellipses demarcate 90% confidence ellipses for each replicate (generation 10, deme 6).

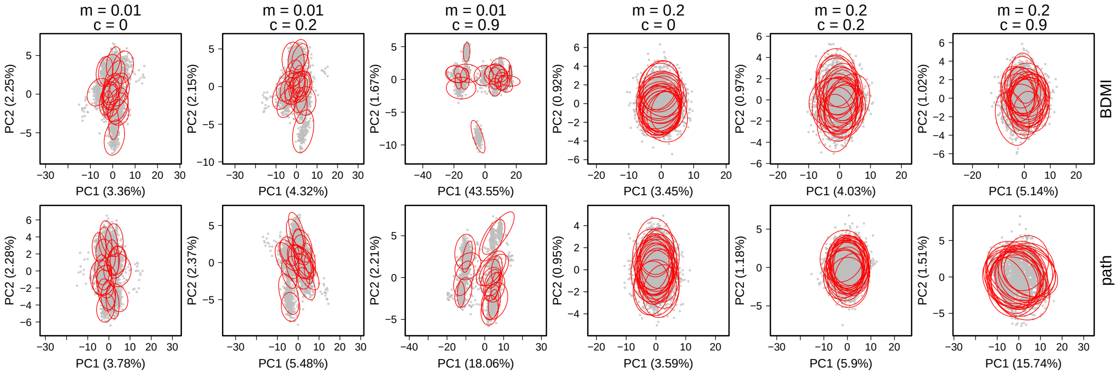

Figure S14: Principal component analysis (PCA) on individual genotypes was used to assess whether replicate simulations differed from one another for a given scenario [migration ($m$); selection ($c$); architecture (BDMI versus pathway); environment (env)]. Gray points correspond to individual scores for the first two PCs, and red ellipses demarcate 90% confidence ellipses for each replicate (generation 100, deme 6).

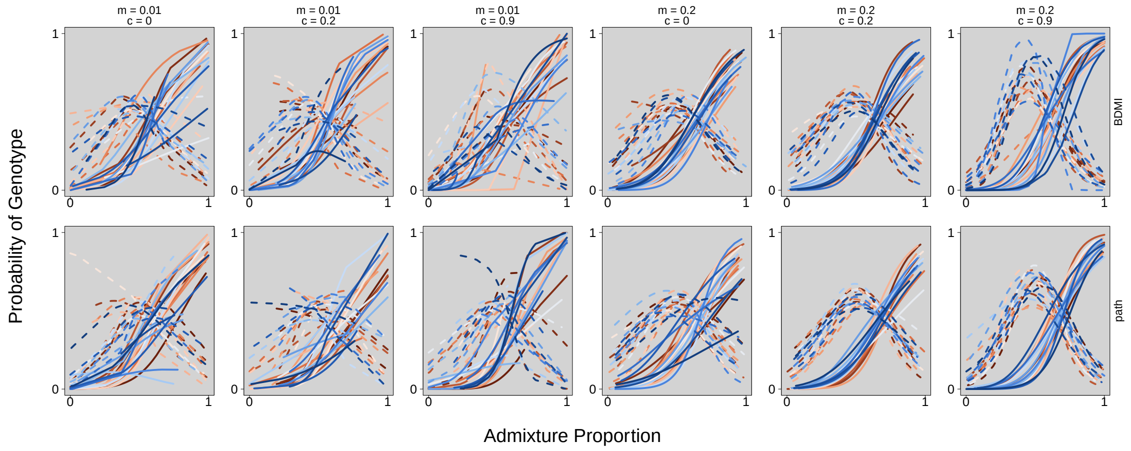

Figure S15: Genomic clines results are shown for the BDMI locus (1.4) across simulated scenarios that differed in levels of gene flow ($m$), selection ($c$), and genetic architecture (BDMI versus pathway). Solid lines show the probability of an AA genotype for a given $q$, whereas dashed lines show the probability of an Aa genotype for a given $q$. Clines are colored by replicate, and the black line represents the mean from all 20 replicates.

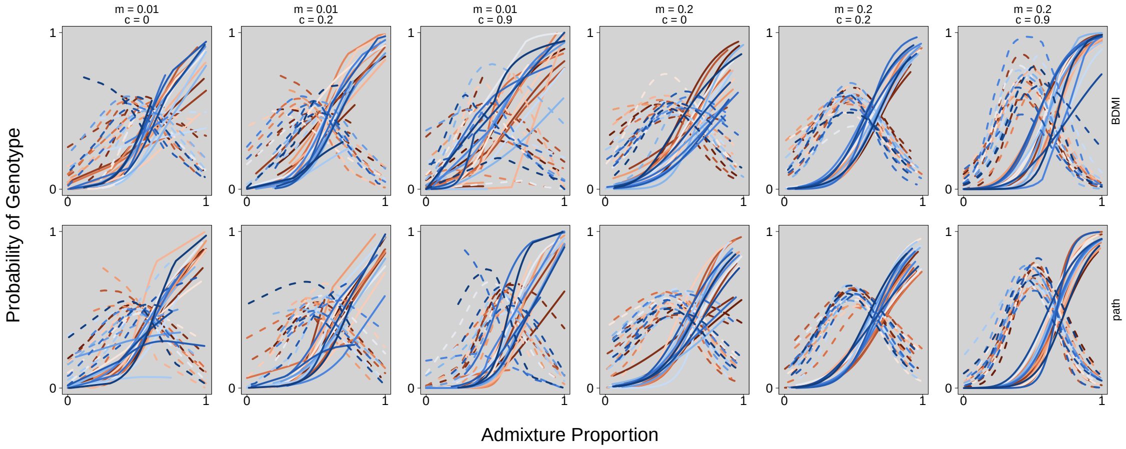

Figure S16: Genomic clines results are shown for the locus residing on the same chromosome as the BDMI (1.10) across simulated scenarios that differed in levels of gene flow ($m$), selection ($c$), and genetic architecture (BDMI versus pathway). Solid lines show the probability of an AA genotype for a given $q$, whereas dashed lines show the probability of an Aa genotype for a given $q$. Clines are colored by replicate, and the black line represents the mean from all 20 replicates.

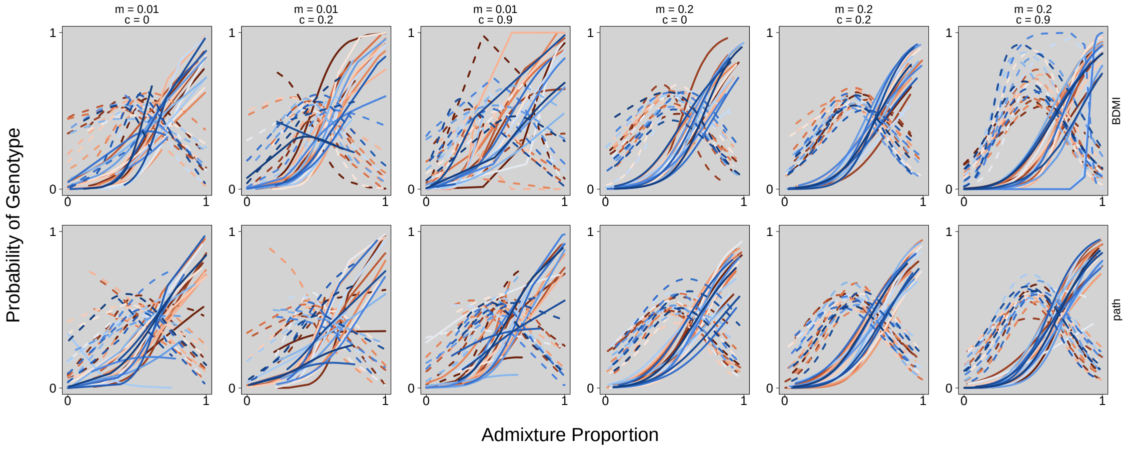

Figure S17: Genomic clines results are shown for the locus residing on a different chromosome than the BDMI (3.4) across simulated scenarios that differed in levels of gene flow ($m$), selection ($c$), and genetic architecture (BDMI versus pathway). Solid lines show the probability of an AA genotype for a given $q$, whereas dashed lines show the probability of an Aa genotype for a given $q$. Clines are colored by replicate, and the black line represents the mean from all 20 replicates.

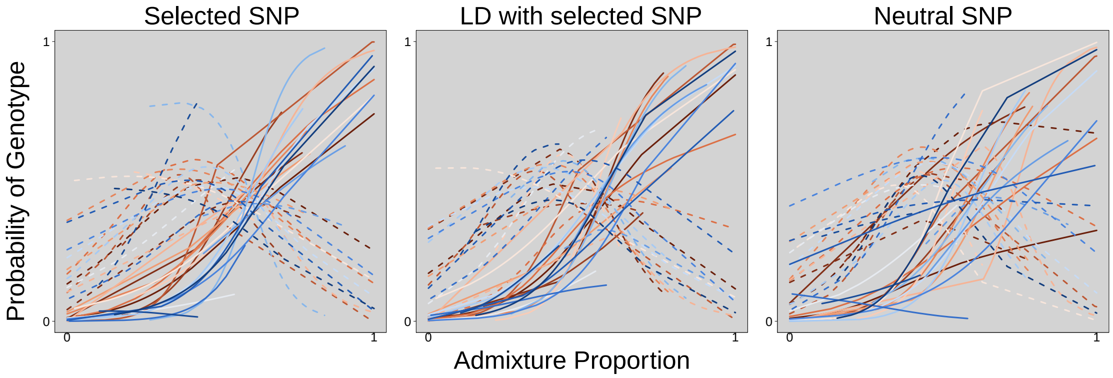

Figure S18: Probability of each AA (homozygous; solid line) and Aa (codominant heterozygote; dashed line) for each of three simulated SNPs in the BDMI scenario where migration is low ($m$=0.01), and selection is high ($c$=0.9), in a 3 deme system. Colors correspond to different replicates and match those in other figures. This multinomial regression shows the probability of either a homozygous or heterozygous genotype for a given admixture proportion $q$.

| Architecture | $m$ | $c$ | Gen | $q$ F | $q$ P | $q$ $R^2$ | $Q$ F | $Q$ P | $Q$ $R^2$ | Junctions $F$ | Junctions $P$ | Junctions $R^2$ |
|---|---|---|---|---|---|---|---|---|---|---|---|---|
| BDMI | 0.01 | 0 | 10 | 35.368 | < 0.001 | 0.184 | 1.845 | 0.014 | 0.012 | 7.024 | < 0.001 | 0.043 |
| BDMI | 0.01 | 0.2 | 10 | 27.457 | < 0.001 | 0.149 | 1.686 | 0.032 | 0.011 | 7.266 | < 0.001 | 0.044 |
| BDMI | 0.01 | 0.9 | 10 | 1148.088 | < 0.001 | 0.880 | 153.268 | < 0.001 | 0.494 | 145.877 | < 0.001 | 0.482 |
| BDMI | 0.2 | 0 | 10 | 3.715 | < 0.001 | 0.023 | 0.820 | 0.685 | 0.005 | 6.924 | < 0.001 | 0.042 |
| BDMI | 0.2 | 0.2 | 10 | 2.160 | < 0.001 | 0.014 | 0.817 | 0.688 | 0.005 | 4.048 | < 0.001 | 0.025 |
| BDMI | 0.2 | 0.9 | 10 | 79.335 | < 0.001 | 0.336 | 16.014 | < 0.001 | 0.093 | 26.778 | < 0.001 | 0.146 |
| Path | 0.01 | 0 | 10 | 51.175 | < 0.001 | 0.246 | 2.006 | 0.006 | 0.013 | 7.336 | < 0.001 | 0.045 |
| Path | 0.01 | 0.2 | 10 | 36.043 | < 0.001 | 0.187 | 1.609 | 0.046 | 0.010 | 4.399 | < 0.001 | 0.027 |
| Path | 0.01 | 0.9 | 10 | 121.459 | < 0.001 | 0.436 | 16.689 | < 0.001 | 0.096 | 11.384 | < 0.001 | 0.068 |
| Path | 0.2 | 0 | 10 | 6.395 | < 0.001 | 0.039 | 0.715 | 0.807 | 0.005 | 3.545 | < 0.001 | 0.022 |
| Path | 0.2 | 0.2 | 10 | 6.935 | < 0.001 | 0.042 | 0.936 | 0.537 | 0.006 | 4.542 | < 0.001 | 0.028 |
| Path | 0.2 | 0.9 | 10 | 7.955 | < 0.001 | 0.048 | 1.066 | 0.380 | 0.007 | 5.254 | < 0.001 | 0.032 |
| BDMI | 0.01 | 0 | 100 | 54.348 | < 0.001 | 0.257 | 7.118 | < 0.001 | 0.043 | 41.355 | < 0.001 | 0.209 |
| BDMI | 0.01 | 0.2 | 100 | 119.427 | < 0.001 | 0.432 | 5.651 | < 0.001 | 0.035 | 40.459 | < 0.001 | 0.205 |
| BDMI | 0.01 | 0.9 | 100 | 3094.124 | < 0.001 | 0.952 | 433.091 | < 0.001 | 0.734 | 1183.061 | < 0.001 | 0.883 |
| BDMI | 0.2 | 0 | 100 | 7.241 | < 0.001 | 0.044 | 2.010 | 0.006 | 0.013 | 21.355 | < 0.001 | 0.120 |
| BDMI | 0.2 | 0.2 | 100 | 25.103 | < 0.001 | 0.138 | 2.217 | 0.002 | 0.014 | 22.976 | < 0.001 | 0.128 |
| BDMI | 0.2 | 0.9 | 100 | 41.213 | < 0.001 | 0.208 | 2.963 | < 0.001 | 0.019 | 44.250 | < 0.001 | 0.220 |
| Path | 0.01 | 0 | 100 | 127.101 | < 0.001 | 0.448 | 4.224 | < 0.001 | 0.026 | 84.769 | < 0.001 | 0.351 |
| Path | 0.01 | 0.2 | 100 | 129.376 | < 0.001 | 0.452 | 11.797 | < 0.001 | 0.070 | 48.581 | < 0.001 | 0.236 |
| Path | 0.01 | 0.9 | 100 | 716.062 | < 0.001 | 0.820 | 22.567 | < 0.001 | 0.126 | 71.015 | < 0.001 | 0.312 |
| Path | 0.2 | 0 | 100 | 12.382 | < 0.001 | 0.073 | 1.628 | 0.042 | 0.010 | 17.761 | < 0.001 | 0.102 |
| Path | 0.2 | 0.2 | 100 | 13.677 | < 0.001 | 0.080 | 1.073 | 0.372 | 0.007 | 30.410 | < 0.001 | 0.162 |
| Path | 0.2 | 0.9 | 100 | 29.223 | < 0.001 | 0.157 | 2.478 | < 0.001 | 0.016 | 30.875 | < 0.001 | 0.164 |

Table S1: ANOVA F statistics were used as a measure of variability among replicate hybrid zones for each of the 12 simulation scenarios [migration ($m$); selection ($c$); architecture (dmi vs. pathway); environment (env)]. Models were run using deme 6 (of 11) for generations 10 or 100, with replicate predicting either admixture proportion ($q$), interspecific ancestry ($Q_{12}$), or the number of junctions.

| Architecture | $m$ | $c$ | Gen | $q$ F | $q$ P | $q$ $R^2$ | $Q$ F | $Q$ P | $Q$ $R^2$ | Junctions $F$ | Junctions $P$ | Junctions $R^2$ |
|---|---|---|---|---|---|---|---|---|---|---|---|---|
| BDMI | 0.01 | 0 | 10 | 21.819 | <0.001 | 0.122 | 4.630 | <0.001 | 0.029 | 5.314 | <0.001 | 0.033 |
| BDMI | 0.01 | 0.2 | 10 | 56.129 | <0.001 | 0.264 | 5.562 | <0.001 | 0.034 | 7.614 | <0.001 | 0.046 |
| BDMI | 0.01 | 0.9 | 10 | 287.737 | <0.001 | 0.647 | 43.040 | <0.001 | 0.215 | 26.256 | <0.001 | 0.143 |
| BDMI | 0.2 | 0 | 10 | 23.371 | <0.001 | 0.130 | 21.926 | <0.001 | 0.123 | 20.648 | <0.001 | 0.116 |
| BDMI | 0.2 | 0.2 | 10 | 10.143 | <0.001 | 0.061 | 9.114 | <0.001 | 0.055 | 9.551 | <0.001 | 0.057 |
| BDMI | 0.2 | 0.9 | 10 | 34.036 | <0.001 | 0.178 | 34.282 | <0.001 | 0.179 | 30.406 | <0.001 | 0.162 |
| Path | 0.01 | 0 | 10 | 23.855 | <0.001 | 0.132 | 1.617 | 0.044 | 0.010 | 5.686 | <0.001 | 0.035 |
| Path | 0.01 | 0.2 | 10 | 44.256 | <0.001 | 0.220 | 4.608 | <0.001 | 0.029 | 6.382 | <0.001 | 0.039 |
| Path | 0.01 | 0.9 | 10 | 22.421 | <0.001 | 0.125 | 2.168 | 0.002 | 0.014 | 5.770 | <0.001 | 0.035 |
| Path | 0.2 | 0 | 10 | 18.745 | <0.001 | 0.107 | 16.313 | <0.001 | 0.094 | 14.213 | <0.001 | 0.083 |
| Path | 0.2 | 0.2 | 10 | 8.675 | <0.001 | 0.052 | 8.353 | <0.001 | 0.051 | 6.897 | <0.001 | 0.042 |
| Path | 0.2 | 0.9 | 10 | 9.957 | <0.001 | 0.060 | 9.841 | <0.001 | 0.059 | 10.189 | <0.001 | 0.061 |
| BDMI | 0.01 | 0 | 100 | 140.951 | <0.001 | 0.473 | 117.390 | <0.001 | 0.428 | 87.023 | <0.001 | 0.357 |
| BDMI | 0.01 | 0.2 | 100 | 273.082 | <0.001 | 0.635 | 198.847 | <0.001 | 0.559 | 137.422 | <0.001 | 0.467 |
| BDMI | 0.01 | 0.9 | 100 | 1001.212 | <0.001 | 0.865 | 699.509 | <0.001 | 0.817 | 384.262 | <0.001 | 0.710 |
| BDMI | 0.2 | 0 | 100 | 5.805 | <0.001 | 0.036 | 5.652 | <0.001 | 0.035 | 43.488 | <0.001 | 0.217 |
| BDMI | 0.2 | 0.2 | 100 | 18.909 | <0.001 | 0.108 | 18.894 | <0.001 | 0.108 | 15.298 | <0.001 | 0.089 |
| BDMI | 0.2 | 0.9 | 100 | 6.420 | <0.001 | 0.039 | 6.387 | <0.001 | 0.039 | 6.125 | <0.001 | 0.038 |
| Path | 0.01 | 0 | 100 | 126.534 | <0.001 | 0.447 | 77.310 | <0.001 | 0.330 | 67.227 | <0.001 | 0.300 |
| Path | 0.01 | 0.2 | 100 | 135.739 | <0.001 | 0.464 | 113.766 | <0.001 | 0.420 | 83.679 | <0.001 | 0.348 |
| Path | 0.01 | 0.9 | 100 | 167.874 | <0.001 | 0.517 | 126.137 | <0.001 | 0.446 | 112.523 | <0.001 | 0.418 |
| Path | 0.2 | 0 | 100 | 2.646 | <0.001 | 0.017 | 2.571 | <0.001 | 0.016 | 32.554 | <0.001 | 0.172 |
| Path | 0.2 | 0.2 | 100 | 4.128 | <0.001 | 0.026 | 4.137 | <0.001 | 0.026 | 26.162 | <0.001 | 0.143 |
| Path | 0.2 | 0.9 | 100 | 4.437 | <0.001 | 0.028 | 4.398 | <0.001 | 0.027 | 35.042 | <0.001 | 0.183 |

Table S2: ANOVA F statistics were used as a measure of variability among replicate hybrid zones for each of the 12 simulation scenarios when one parental population was fifty times as large as the other parental population [migration ($m$); selection ($c$); architecture (dmi vs. pathway); environment (env)]. Models were run using deme 2 (of 3) for generations 10 or 100, with replicate predicting either admixture proportion ($q$), interspecific ancestry ($Q_{12}$), or the number of junctions.

| Architecture | $m$ | $c$ | Gen | $q$ F | $q$ P | $q$ $R^2$ | $Q$ F | $Q$ P | $Q$ $R^2$ | Junctions $F$ | Junctions $P$ | Junctions $R^2$ |
|---|---|---|---|---|---|---|---|---|---|---|---|---|
| BDMI | 0.01 | 0 | 10 | 15.748 | <0.001 | 0.091 | 1.844 | 0.014 | 0.012 | 5.549 | <0.001 | 0.034 |
| BDMI | 0.01 | 0.2 | 10 | 25.698 | <0.001 | 0.141 | 2.474 | <0.001 | 0.016 | 4.612 | <0.001 | 0.029 |
| BDMI | 0.01 | 0.9 | 10 | 213.724 | <0.001 | 0.577 | 79.131 | <0.001 | 0.335 | 41.310 | <0.001 | 0.208 |
| BDMI | 0.2 | 0 | 10 | 9.705 | <0.001 | 0.058 | 8.769 | <0.001 | 0.053 | 14.218 | <0.001 | 0.083 |
| BDMI | 0.2 | 0.2 | 10 | 8.341 | <0.001 | 0.050 | 8.795 | <0.001 | 0.053 | 8.429 | <0.001 | 0.051 |
| BDMI | 0.2 | 0.9 | 10 | 3.883 | <0.001 | 0.024 | 5.559 | <0.001 | 0.034 | 10.755 | <0.001 | 0.064 |
| Path | 0.01 | 0 | 10 | 35.244 | <0.001 | 0.183 | 3.728 | <0.001 | 0.023 | 3.922 | <0.001 | 0.024 |
| Path | 0.01 | 0.2 | 10 | 36.794 | <0.001 | 0.190 | 1.688 | 0.031 | 0.011 | 10.376 | <0.001 | 0.062 |
| Path | 0.01 | 0.9 | 10 | 19.534 | <0.001 | 0.111 | 2.549 | <0.001 | 0.016 | 5.946 | <0.001 | 0.037 |
| Path | 0.2 | 0 | 10 | 7.225 | <0.001 | 0.044 | 7.982 | <0.001 | 0.048 | 10.118 | <0.001 | 0.061 |
| Path | 0.2 | 0.2 | 10 | 4.874 | <0.001 | 0.030 | 4.866 | <0.001 | 0.030 | 5.385 | <0.001 | 0.033 |
| Path | 0.2 | 0.9 | 10 | 6.415 | <0.001 | 0.039 | 7.001 | <0.001 | 0.043 | 8.676 | <0.001 | 0.052 |
| BDMI | 0.01 | 0 | 100 | 109.085 | <0.001 | 0.410 | 85.468 | <0.001 | 0.353 | 79.943 | <0.001 | 0.338 |
| BDMI | 0.01 | 0.2 | 100 | 109.594 | <0.001 | 0.411 | 70.923 | <0.001 | 0.311 | 79.627 | <0.001 | 0.337 |
| BDMI | 0.01 | 0.9 | 100 | 359.042 | <0.001 | 0.696 | 342.774 | <0.001 | 0.686 | 289.251 | <0.001 | 0.648 |
| BDMI | 0.2 | 0 | 100 | 6.988 | <0.001 | 0.043 | 7.752 | <0.001 | 0.047 | 17.725 | <0.001 | 0.102 |
| BDMI | 0.2 | 0.2 | 100 | 2.774 | <0.001 | 0.017 | 2.814 | <0.001 | 0.018 | 16.161 | <0.001 | 0.093 |
| BDMI | 0.2 | 0.9 | 100 | 2.708 | <0.001 | 0.017 | 4.578 | <0.001 | 0.028 | 16.898 | <0.001 | 0.097 |
| Path | 0.01 | 0 | 100 | 148.320 | <0.001 | 0.486 | 99.982 | <0.001 | 0.389 | 76.859 | <0.001 | 0.329 |
| Path | 0.01 | 0.2 | 100 | 106.111 | <0.001 | 0.404 | 85.729 | <0.001 | 0.353 | 50.050 | <0.001 | 0.242 |
| Path | 0.01 | 0.9 | 100 | 59.603 | <0.001 | 0.275 | 39.986 | <0.001 | 0.203 | 121.410 | <0.001 | 0.436 |
| Path | 0.2 | 0 | 100 | 5.226 | <0.001 | 0.032 | 5.695 | <0.001 | 0.035 | 30.241 | <0.001 | 0.162 |
| Path | 0.2 | 0.2 | 100 | 2.332 | 0.001 | 0.015 | 2.764 | <0.001 | 0.017 | 7.111 | <0.001 | 0.043 |
| Path | 0.2 | 0.9 | 100 | 5.257 | <0.001 | 0.032 | 5.951 | <0.001 | 0.037 | 19.513 | <0.001 | 0.111 |

Table S3: ANOVA F statistics were used as a measure of variability among replicate hybrid zones for each of the 12 simulation scenarios when one parental population was ten times as large as the other parental population [migration ($m$); selection ($c$); architecture (dmi vs. pathway); environment (env)]. Models were run using deme 2 (of 3) for generations 10 or 100, with replicate predicting either admixture proportion ($q$), interspecific ancestry ($Q_{12}$), or the number of junctions.

| Architecture | $m$ | $c$ | Gen | $q$ F | $q$ P | $q$ $R^2$ | $Q$ F | $Q$ P | $Q$ $R^2$ | Junctions $F$ | Junctions $P$ | Junctions $R^2$ |
|---|---|---|---|---|---|---|---|---|---|---|---|---|
| BDMI | 0.01 | 0 | 10 | 23.387 | <0.001 | 0.130 | 0.705 | 0.818 | 0.004 | 2.955 | <0.001 | 0.018 |
| BDMI | 0.01 | 0.2 | 10 | 37.560 | <0.001 | 0.193 | 2.657 | <0.001 | 0.017 | 7.147 | <0.001 | 0.044 |
| BDMI | 0.01 | 0.9 | 10 | 481.211 | <0.001 | 0.754 | 121.935 | <0.001 | 0.437 | 92.774 | <0.001 | 0.372 |
| BDMI | 0.2 | 0 | 10 | 2.240 | 0.002 | 0.014 | 1.653 | 0.037 | 0.010 | 5.473 | <0.001 | 0.034 |
| BDMI | 0.2 | 0.2 | 10 | 4.070 | <0.001 | 0.025 | 5.164 | <0.001 | 0.032 | 8.309 | <0.001 | 0.050 |
| BDMI | 0.2 | 0.9 | 10 | 1.977 | 0.007 | 0.012 | 4.768 | <0.001 | 0.030 | 13.036 | <0.001 | 0.077 |
| Path | 0.01 | 0 | 10 | 49.576 | <0.001 | 0.240 | 2.697 | <0.001 | 0.017 | 4.410 | <0.001 | 0.027 |
| Path | 0.01 | 0.2 | 10 | 23.114 | <0.001 | 0.128 | 3.329 | <0.001 | 0.021 | 4.903 | <0.001 | 0.030 |
| Path | 0.01 | 0.9 | 10 | 35.223 | <0.001 | 0.183 | 2.630 | <0.001 | 0.016 | 8.165 | <0.001 | 0.049 |
| Path | 0.2 | 0 | 10 | 8.345 | <0.001 | 0.051 | 8.708 | <0.001 | 0.053 | 15.038 | <0.001 | 0.087 |
| Path | 0.2 | 0.2 | 10 | 3.927 | <0.001 | 0.024 | 4.764 | <0.001 | 0.029 | 4.251 | <0.001 | 0.026 |
| Path | 0.2 | 0.9 | 10 | 2.743 | <0.001 | 0.017 | 3.092 | <0.001 | 0.019 | 4.118 | <0.001 | 0.026 |
| BDMI | 0.01 | 0 | 100 | 131.103 | <0.001 | 0.455 | 98.103 | <0.001 | 0.385 | 136.322 | <0.001 | 0.465 |
| BDMI | 0.01 | 0.2 | 100 | 80.531 | <0.001 | 0.339 | 61.135 | <0.001 | 0.280 | 63.959 | <0.001 | 0.290 |
| BDMI | 0.01 | 0.9 | 100 | 348.977 | <0.001 | 0.690 | 348.465 | <0.001 | 0.690 | 298.815 | <0.001 | 0.656 |
| BDMI | 0.2 | 0 | 100 | 7.174 | <0.001 | 0.044 | 6.968 | <0.001 | 0.043 | 18.690 | <0.001 | 0.106 |
| BDMI | 0.2 | 0.2 | 100 | 5.777 | <0.001 | 0.036 | 7.094 | <0.001 | 0.043 | 13.734 | <0.001 | 0.081 |
| BDMI | 0.2 | 0.9 | 100 | 2.484 | <0.001 | 0.016 | 3.715 | <0.001 | 0.023 | 10.828 | <0.001 | 0.065 |
| Path | 0.01 | 0 | 100 | 96.659 | <0.001 | 0.381 | 64.298 | <0.001 | 0.291 | 101.029 | <0.001 | 0.392 |
| Path | 0.01 | 0.2 | 100 | 44.461 | <0.001 | 0.221 | 39.108 | <0.001 | 0.200 | 57.741 | <0.001 | 0.269 |
| Path | 0.01 | 0.9 | 100 | 43.654 | <0.001 | 0.218 | 33.062 | <0.001 | 0.174 | 74.611 | <0.001 | 0.322 |
| Path | 0.2 | 0 | 100 | 4.817 | <0.001 | 0.030 | 5.200 | <0.001 | 0.032 | 11.105 | <0.001 | 0.066 |
| Path | 0.2 | 0.2 | 100 | 5.224 | <0.001 | 0.032 | 7.471 | <0.001 | 0.045 | 24.587 | <0.001 | 0.136 |
| Path | 0.2 | 0.9 | 100 | 2.346 | 0.001 | 0.015 | 3.905 | <0.001 | 0.024 | 6.958 | <0.001 | 0.042 |

Table S4: ANOVA F statistics were used as a measure of variability among replicate hybrid zones for each of the 12 simulation scenarios when one parental population was five times as large as the other parental population [migration ($m$); selection ($c$); architecture (dmi vs. pathway); environment (env)]. Models were run using deme 2 (of 3) for generations 10 or 100, with replicate predicting either admixture proportion ($q$), interspecific ancestry ($Q_{12}$), or the number of junctions.

| Architecture | $m$ | $c$ | Gen | $q$ F | $q$ P | $q$ $R^2$ | $Q$ F | $Q$ P | $Q$ $R^2$ | Junctions $F$ | Junctions $P$ | Junctions $R^2$ |
|---|---|---|---|---|---|---|---|---|---|---|---|---|
| BDMI | 0.01 | 0 | 10 | 51.903 | <0.001 | 0.249 | 2.126 | 0.003 | 0.013 | 7.163 | <0.001 | 0.044 |
| BDMI | 0.01 | 0.2 | 10 | 37.153 | <0.001 | 0.192 | 1.706 | 0.029 | 0.011 | 7.028 | <0.001 | 0.043 |
| BDMI | 0.01 | 0.9 | 10 | 180.083 | <0.001 | 0.534 | 52.171 | <0.001 | 0.250 | 42.227 | <0.001 | 0.212 |
| BDMI | 0.2 | 0 | 10 | 3.214 | <0.001 | 0.020 | 2.958 | <0.001 | 0.019 | 4.300 | <0.001 | 0.027 |
| BDMI | 0.2 | 0.2 | 10 | 2.755 | <0.001 | 0.017 | 2.777 | <0.001 | 0.017 | 5.578 | <0.001 | 0.034 |
| BDMI | 0.2 | 0.9 | 10 | 1.667 | 0.035 | 0.011 | 3.603 | <0.001 | 0.022 | 6.088 | <0.001 | 0.037 |
| Path | 0.01 | 0 | 10 | 14.715 | <0.001 | 0.086 | 0.900 | 0.583 | 0.006 | 6.427 | <0.001 | 0.039 |
| Path | 0.01 | 0.2 | 10 | 33.461 | <0.001 | 0.176 | 2.915 | <0.001 | 0.018 | 8.436 | <0.001 | 0.051 |
| Path | 0.01 | 0.9 | 10 | 22.898 | <0.001 | 0.127 | 2.565 | <0.001 | 0.016 | 5.867 | <0.001 | 0.036 |
| Path | 0.2 | 0 | 10 | 1.594 | 0.049 | 0.010 | 2.065 | 0.004 | 0.013 | 4.882 | <0.001 | 0.030 |
| Path | 0.2 | 0.2 | 10 | 1.405 | 0.113 | 0.009 | 1.294 | 0.176 | 0.008 | 3.868 | <0.001 | 0.024 |
| Path | 0.2 | 0.9 | 10 | 2.000 | 0.006 | 0.013 | 2.076 | 0.004 | 0.013 | 1.703 | 0.029 | 0.011 |
| BDMI | 0.01 | 0 | 100 | 64.942 | <0.001 | 0.293 | 26.523 | <0.001 | 0.145 | 33.337 | <0.001 | 0.175 |
| BDMI | 0.01 | 0.2 | 100 | 35.450 | <0.001 | 0.184 | 15.335 | <0.001 | 0.089 | 53.818 | <0.001 | 0.255 |
| BDMI | 0.01 | 0.9 | 100 | 55.107 | <0.001 | 0.260 | 71.967 | <0.001 | 0.315 | 126.356 | <0.001 | 0.446 |
| BDMI | 0.2 | 0 | 100 | 2.161 | 0.002 | 0.014 | 3.088 | <0.001 | 0.019 | 6.491 | <0.001 | 0.040 |
| BDMI | 0.2 | 0.2 | 100 | 3.746 | <0.001 | 0.023 | 2.139 | 0.003 | 0.013 | 3.775 | <0.001 | 0.024 |
| BDMI | 0.2 | 0.9 | 100 | 1.160 | 0.283 | 0.007 | 2.249 | 0.001 | 0.014 | 5.786 | <0.001 | 0.036 |
| Path | 0.01 | 0 | 100 | 46.144 | <0.001 | 0.227 | 29.592 | <0.001 | 0.159 | 32.953 | <0.001 | 0.174 |
| Path | 0.01 | 0.2 | 100 | 45.305 | <0.001 | 0.224 | 29.134 | <0.001 | 0.157 | 39.003 | <0.001 | 0.199 |
| Path | 0.01 | 0.9 | 100 | 20.442 | <0.001 | 0.115 | 12.873 | <0.001 | 0.076 | 30.556 | <0.001 | 0.163 |
| Path | 0.2 | 0 | 100 | 3.175 | <0.001 | 0.020 | 4.318 | <0.001 | 0.027 | 9.719 | <0.001 | 0.058 |
| Path | 0.2 | 0.2 | 100 | 3.523 | <0.001 | 0.022 | 2.180 | 0.002 | 0.014 | 4.294 | <0.001 | 0.027 |
| Path | 0.2 | 0.9 | 100 | 1.402 | 0.114 | 0.009 | 1.475 | 0.084 | 0.009 | 4.228 | <0.001 | 0.026 |

Table S5: ANOVA F statistics were used as a measure of variability among replicate hybrid zones for each of the 12 simulation scenarios when one parental population was twice as large as the other parental population [migration ($m$); selection ($c$); architecture (dmi vs. pathway); environment (env)]. Models were run using deme 2 (of 3) for generations 10 or 100, with replicate predicting either admixture proportion ($q$), interspecific ancestry ($Q_{12}$), or the number of junctions.

| Architecture | $m$ | $c$ | Gen | $q$ F | $q$ P | $q$ $R^2$ | $Q$ F | $Q$ P | $Q$ $R^2$ | Junctions $F$ | Junctions $P$ | Junctions $R^2$ |
|---|---|---|---|---|---|---|---|---|---|---|---|---|
| BDMI | 0.01 | 0 | 10 | 23.263 | <0.001 | 0.129 | 1.627 | 0.042 | 0.010 | 7.248 | <0.001 | 0.044 |
| BDMI | 0.01 | 0.2 | 10 | 38.828 | <0.001 | 0.198 | 1.543 | 0.062 | 0.010 | 3.393 | <0.001 | 0.021 |
| BDMI | 0.01 | 0.9 | 10 | 395.569 | <0.001 | 0.716 | 49.665 | <0.001 | 0.241 | 38.403 | <0.001 | 0.197 |
| BDMI | 0.2 | 0 | 10 | 3.122 | <0.001 | 0.013 | 2.060 | 0.004 | 0.009 | 3.276 | <0.001 | 0.014 |
| BDMI | 0.2 | 0.2 | 10 | 4.170 | <0.001 | 0.026 | 1.362 | 0.135 | 0.009 | 5.420 | <0.001 | 0.033 |
| BDMI | 0.2 | 0.9 | 10 | 6.382 | <0.001 | 0.039 | 7.882 | <0.001 | 0.048 | 11.696 | <0.001 | 0.069 |
| Path | 0.01 | 0 | 10 | 35.034 | <0.001 | 0.183 | 2.071 | 0.004 | 0.013 | 5.263 | <0.001 | 0.032 |
| Path | 0.01 | 0.2 | 10 | 53.467 | <0.001 | 0.254 | 2.439 | <0.001 | 0.015 | 6.012 | <0.001 | 0.037 |
| Path | 0.01 | 0.9 | 10 | 35.299 | <0.001 | 0.184 | 1.323 | 0.157 | 0.008 | 5.908 | <0.001 | 0.036 |
| Path | 0.2 | 0 | 10 | 2.221 | 0.002 | 0.014 | 1.739 | 0.024 | 0.011 | 3.217 | <0.001 | 0.020 |
| Path | 0.2 | 0.2 | 10 | 6.732 | <0.001 | 0.041 | 1.248 | 0.208 | 0.008 | 4.226 | <0.001 | 0.026 |
| Path | 0.2 | 0.9 | 10 | 4.955 | <0.001 | 0.031 | 1.350 | 0.141 | 0.009 | 3.427 | <0.001 | 0.021 |
| BDMI | 0.01 | 0 | 100 | 55.981 | <0.001 | 0.263 | 7.904 | <0.001 | 0.048 | 36.488 | <0.001 | 0.189 |
| BDMI | 0.01 | 0.2 | 100 | 49.674 | <0.001 | 0.241 | 6.327 | <0.001 | 0.039 | 18.442 | <0.001 | 0.105 |
| BDMI | 0.01 | 0.9 | 100 | 989.668 | <0.001 | 0.863 | 249.315 | <0.001 | 0.614 | 213.865 | <0.001 | 0.577 |
| BDMI | 0.2 | 0 | 100 | 1.284 | 0.182 | 0.005 | 0.423 | 0.986 | 0.002 | 1.241 | 0.213 | 0.005 |
| BDMI | 0.2 | 0.2 | 100 | 4.288 | <0.001 | 0.027 | 0.984 | 0.477 | 0.006 | 10.601 | <0.001 | 0.063 |
| BDMI | 0.2 | 0.9 | 100 | 2.831 | <0.001 | 0.018 | 1.910 | 0.010 | 0.012 | 6.313 | <0.001 | 0.039 |
| Path | 0.01 | 0 | 100 | 67.013 | <0.001 | 0.299 | 7.261 | <0.001 | 0.044 | 43.262 | <0.001 | 0.216 |
| Path | 0.01 | 0.2 | 100 | 53.636 | <0.001 | 0.255 | 7.709 | <0.001 | 0.047 | 31.164 | <0.001 | 0.166 |
| Path | 0.01 | 0.9 | 100 | 67.021 | <0.001 | 0.299 | 18.403 | <0.001 | 0.105 | 55.266 | <0.001 | 0.261 |
| Path | 0.2 | 0 | 100 | 3.407 | <0.001 | 0.021 | 1.141 | 0.301 | 0.007 | 4.466 | <0.001 | 0.028 |
| Path | 0.2 | 0.2 | 100 | 2.981 | <0.001 | 0.019 | 0.908 | 0.573 | 0.006 | 16.286 | <0.001 | 0.094 |
| Path | 0.2 | 0.9 | 100 | 2.683 | <0.001 | 0.017 | 0.619 | 0.895 | 0.004 | 7.351 | <0.001 | 0.045 |

Table S6: ANOVA F statistics were used as a measure of variability among replicate hybrid zones for each of the 12 simulation scenarios when parental populations were the same size [migration ($m$); selection ($c$); architecture (dmi vs. pathway); environment (env)]. Models were run using deme 2 (of 3) for generations 10 or 100, with replicate predicting either admixture proportion ($q$), interspecific ancestry ($Q_{12}$), or the number of junctions.

| Architecture | $m$ | $c$ | Gen | loc 1.4 F | loc 1.4 P | loc 1.4 $R^2$ | loc 1.10 F | loc 1.10 P | loc 1.10 $R^2$ | loc 3.4 F | loc 3.4 P | loc 3.4 $R^2$ |
|---|---|---|---|---|---|---|---|---|---|---|---|---|
| BDMI | 0.01 | 0 | 10 | 14.782 | 0.000 | 0.086 | 9.711 | 0.000 | 0.058 | 6.943 | 0.000 | 0.042 |
| BDMI | 0.01 | 0.2 | 10 | 8.186 | 0.000 | 0.050 | 14.947 | 0.000 | 0.087 | 8.894 | 0.000 | 0.054 |
| BDMI | 0.01 | 0.9 | 10 | 233.132 | 0.000 | 0.598 | 280.900 | 0.000 | 0.642 | 134.425 | 0.000 | 0.462 |
| BDMI | 0.2 | 0 | 10 | 5.279 | 0.000 | 0.033 | 5.058 | 0.000 | 0.031 | 3.385 | 0.000 | 0.021 |
| BDMI | 0.2 | 0.2 | 10 | 2.724 | 0.000 | 0.017 | 1.825 | 0.016 | 0.012 | 2.272 | 0.001 | 0.014 |
| BDMI | 0.2 | 0.9 | 10 | 62.338 | 0.000 | 0.284 | 66.285 | 0.000 | 0.297 | 51.634 | 0.000 | 0.248 |
| Path | 0.01 | 0 | 10 | 18.742 | 0.000 | 0.107 | 7.191 | 0.000 | 0.044 | 12.637 | 0.000 | 0.075 |
| Path | 0.01 | 0.2 | 10 | 10.396 | 0.000 | 0.062 | 8.560 | 0.000 | 0.052 | 7.683 | 0.000 | 0.047 |
| Path | 0.01 | 0.9 | 10 | 25.062 | 0.000 | 0.138 | 58.938 | 0.000 | 0.273 | 22.426 | 0.000 | 0.125 |
| Path | 0.2 | 0 | 10 | 6.309 | 0.000 | 0.039 | 4.033 | 0.000 | 0.025 | 3.742 | 0.000 | 0.023 |
| Path | 0.2 | 0.2 | 10 | 4.370 | 0.000 | 0.027 | 2.769 | 0.000 | 0.017 | 6.042 | 0.000 | 0.037 |
| Path | 0.2 | 0.9 | 10 | 6.966 | 0.000 | 0.043 | 7.139 | 0.000 | 0.044 | 4.208 | 0.000 | 0.026 |

Table S7: We used ANOVA to ask whether there was significant variation in the genotypes (0,1,2) among replicates for three focal loci: a locus under selection (1.4); a physically linked locus (1.10); and a physically unlinked locus (3.4). This was done using deme 6 (of 11).

| Architecture | m | c | Gen | loc 1.4 F | loc 1.4 P | loc 1.4 cor | loc 1.10 F | loc 1.10 P | loc 1.10 cor | loc 3.4 F | loc 3.4 P | loc 3.4 cor |
|---|---|---|---|---|---|---|---|---|---|---|---|---|
| BDMI | 0.01 | 0 | 10 | 10.500 | 0.000 | -0.019 | 5.606 | 0.000 | -0.124 | 4.437 | 0.000 | -0.028 |
| BDMI | 0.01 | 0.2 | 10 | 6.710 | 0.000 | -0.012 | 6.723 | 0.000 | -0.031 | 8.807 | 0.000 | -0.126 |
| BDMI | 0.01 | 0.9 | 10 | 25.253 | 0.000 | -0.070 | 24.782 | 0.000 | 0.047 | 29.962 | 0.000 | -0.020 |
| BDMI | 0.2 | 0 | 10 | 1.464 | 0.088 | 0.014 | 2.015 | 0.006 | 0.006 | 2.893 | 0.000 | -0.138 |
| BDMI | 0.2 | 0.2 | 10 | 2.010 | 0.006 | -0.032 | 1.211 | 0.238 | 0.067 | 1.991 | 0.006 | -0.042 |
| BDMI | 0.2 | 0.9 | 10 | 8.596 | 0.000 | 0.207 | 10.045 | 0.000 | -0.010 | 7.248 | 0.000 | -0.143 |
| Path | 0.01 | 0 | 10 | 9.713 | 0.000 | 0.030 | 5.212 | 0.000 | 0.086 | 6.300 | 0.000 | 0.061 |
| Path | 0.01 | 0.2 | 10 | 3.576 | 0.000 | -0.125 | 7.406 | 0.000 | 0.067 | 6.639 | 0.000 | -0.027 |
| Path | 0.01 | 0.9 | 10 | 7.998 | 0.000 | 0.100 | 21.770 | 0.000 | 0.019 | 14.305 | 0.000 | 0.074 |
| Path | 0.2 | 0 | 10 | 3.399 | 0.000 | -0.034 | 2.184 | 0.002 | -0.095 | 2.914 | 0.000 | 0.042 |
| Path | 0.2 | 0.2 | 10 | 2.080 | 0.004 | 0.022 | 1.112 | 0.331 | 0.006 | 3.472 | 0.000 | -0.113 |
| Path | 0.2 | 0.9 | 10 | 2.771 | 0.000 | -0.239 | 2.611 | 0.000 | 0.020 | 1.443 | 0.096 | 0.031 |

Table S8: We used ANOVA to ask whether there was significant variation in the minor ancestries among replicates for three focal loci: a locus under selection (1.4); a physically linked locus (1.10); and a physically unlinked locus (3.4). We also examined the correlation of minor ancestry at each locus between replicate population 1 and population 2, to determine if minor ancestry was consistently depressed across replicates, and found low correlations.
